# A double-ring of human RAD52 remodels replication forks restricting fork reversal

**DOI:** 10.1101/2023.11.14.566657

**Authors:** Masayoshi Honda, Mortezaali Razzaghi, Paras Gaur, Eva Malacaria, Giorgia Marozzi, Ludovica Di Biagi, Francesca Antonella Aiello, Emeleeta A. Paintsil, Andrew J. Stanfield, Bailey J. Deppe, Lokesh Gakhar, Nicholas J. Schnicker, M. Ashley Spies, Pietro Pichierri, Maria Spies

## Abstract

Human RAD52^1,2^ is a multifunctional DNA repair protein involved in several cellular events that support genome stability including protection of stalled DNA replication forks from excessive degradation^3–7^. In its gatekeeper role, RAD52 binds to and stabilizes stalled replication forks during replication stress protecting them from reversal by SMARCAL1^5^. The structural and molecular mechanism of the RAD52-mediated fork protection remains elusive. Here, using P1 nuclease sensitivity, biochemical and single-molecule analyses we show that RAD52 dynamically remodels replication forks through its strand exchange activity. The presence of the ssDNA binding protein RPA at the fork modulates the kinetics of the strand exchange without impeding the reaction outcome. Mass photometry and single-particle cryo-electron microscopy show that the replication fork promotes a unique nucleoprotein structure containing head-to-head arrangement of two undecameric RAD52 rings with an extended positively charged surface that accommodates all three arms of the replication fork. We propose that the formation and continuity of this surface is important for the strand exchange reaction and for competition with SMARCAL1.

**One Sentence Summary:** Using cryo-EM, biochemical and single-molecule approaches we show that the structure of stalled DNA replication fork promotes a unique two-ring organization of human RAD52 protein which remodels the fork via DNA strand exchange.

## Introduction

The accurate and timely DNA replication program is a prerequisite of a stable genome. Some of the stalled replication forks are reversed into so-called “chicken foot” structures^8,9^. If employed indiscriminately, fork reversal may cause excessive fork erosion and genome instability^9^. Among its many cellular functions, human DNA repair protein RAD52^1,2^ is recruited to ssDNA at stalled or damaged DNA replication forks where it antagonizes fork reversal by motor enzymes, such as SMARCAL1^5^ (reviewed in ^3,4,10^). This gatekeeper role of RAD52 is distinct from its residual activity in homologous recombinaiton^11,12^, as well as RAD52 functions in potentially mutagenic single-strand annealing (SSA)^13^, antagonizing DNA polymerase θ and theta-mediated end joining^14^, recruitment of MRE11 nuclease to reversed forks^15^, or mitotic DNA synthesis (MiDAS)^16,17^, but is related to fork cleavage by RAD52/MUS81 axis in response to prolonged replication stress^18^. The RAD52 gatekeeper function is temporally upstream of the reversed fork protection by recombination machinery and may therefore represent a basis for the synthetically lethal interaction between RAD52 loss or pharmacological inhibition and BRCAness^3,6,7,11,19–26^. In the absence of RAD52, or upon RAD52 inhibition, the reversed forks are restored, but with large ssDNA gaps, *i.e.* loss of RAD52 or its inhibition does not result in a loss of both nascent strands at the arrested fork, but degradation of just one of the two nascent strands leaving the remaining nascent ssDNA in a detectable single-stranded form^5^. Notably, persistence of the post-replicative ssDNA gaps was shown to be toxic to BRCA-deficient cells^27^. The molecular and structural mechanisms of the RAD52 action at stalled replication work, and whether the interactions involved are similar to those occurring during strand annealing remained unknown.

The main isoform of human RAD52 is a 418 amino acids long protein comprised by an evolutionarily conserved self-oligomerization N-terminal domain that interacts with DNA and RNA, and a species-specific disordered C-terminal domain involved in protein-protein interactions^3,6^. Biochemical activities associated with the RAD52 include binding of single- and double-stranded DNA, RNA and DNA/RNA hybrids, annealing of complementary DNA strands coated with ssDNA binding protein RPA, and an inverse strand exchange reaction between homologous dsDNA and ssRNA or DNA^28–33^. RAD52 forms a ring-shaped oligomer ranging from heptamer to undecamer^25,34^. Crystal structures of the N-terminal domain showed an undecameric RAD52 ring^32,35,36^. Single-particle cryo-EM structures confirmed that the N-terminal domain of the full-length human RAD52 also forms an undecameric ring, while the C-terminal domain is disordered^37,38^. Open decameric rings of RAD52 were recently observed in the RAD52-ssDNA and RAD52-ssDNA-RPA complexes poised for strand annealing^39^. Despite forming decameric rings, the N-terminal domain of yeast Rad52 displays a high degree of structural similarity with human RAD52^40^.

Previously, we showed that RAD52 binds to the replication fork interacting with all three arms of the three-way junction making the fork refractory to remodeling by SMARCAL1^5^. The mechanism by which this protection is achieved remained unresolved. Here, we show that a synthetic DNA structure mimicking stalled DNA replication forks recruits two undecameric RAD52 rings that are arranged in a head-to-head manner. At the fork, RAD52 facilities a dynamic exchange between homologous ssDNA and dsDNA (even when ssDNA is bound by RPA) converting the fork into a four-way junction. Competition with SMARCAL1, however, is through direct binding and sequestering the three arms of the fork by the two rings of RAD52.

## Results

### Human RAD52 protein targets DNA replication forks and facilitates fork remodeling by exchanging strands between the leading and lagging arms

RAD52 can protect replication forks from unscheduled reversal by two non-mutually exclusive mechanisms. The simplest mechanism involves binding of the RAD52 to both leading and lagging strand thereby preventing SMARCAL1 access. RAD52, however, also has the capacity to carry out so-called inverse strand exchange reaction between homologous double- and single-stranded nucleic acids^33^. A stalled replication fork containing either a leading or a lagging strand gap offers a convenient proximity for the homologous single- and double-stranded arms. To test whether RAD52 can exchange DNA strands at the replication fork we constructed the DNA junction resembling stalled replication fork with a leading strand gap of 30 nucleotides (**Fig.1A**; **Supplemental Table S1**). The parental leading strand is labeled with the Cy3 dye at the 5’-end, while the nascent lagging strand is labeled with the Cy5 dye at the 3’-end. In the absence of RAD52, the Cy5 strand exists in a fully double stranded form and therefore is protected from digestion by the ssDNA specific nuclease P1. In contrast, the ssDNA gap on the Cy3 strand is susceptible to P1 cleavage yielding a distinct 22-nucleotide product. The DNA strand exchange between the two homologous arms of the fork converts the Cy3 strand to a fully duplex structure, while exposing the Cy5 strand. **Fig.1C-E** shows that RAD52 action on the fork DNA containing homologous arms results in the Cy3 strand protection form and the Cy5 strand sensitization to P1 nuclease. The appearance of a distinct 22-nucleotide product in the Cy5 channel indicates that the strand exchange reaction is limited to the ssDNA gap region and does not result in further branch migration into homologous duplex. The maximum Cy5 strand sensitization was achieved at the stoichiometry of approximately two RAD52 undecamers per fork. On the heterologous fork, we observed only the Cy3 strand protection, but no exposure of the Cy5 strand (**Fig. 1 F&G**; quantification in **Fig1E**).

**Figure 1.**
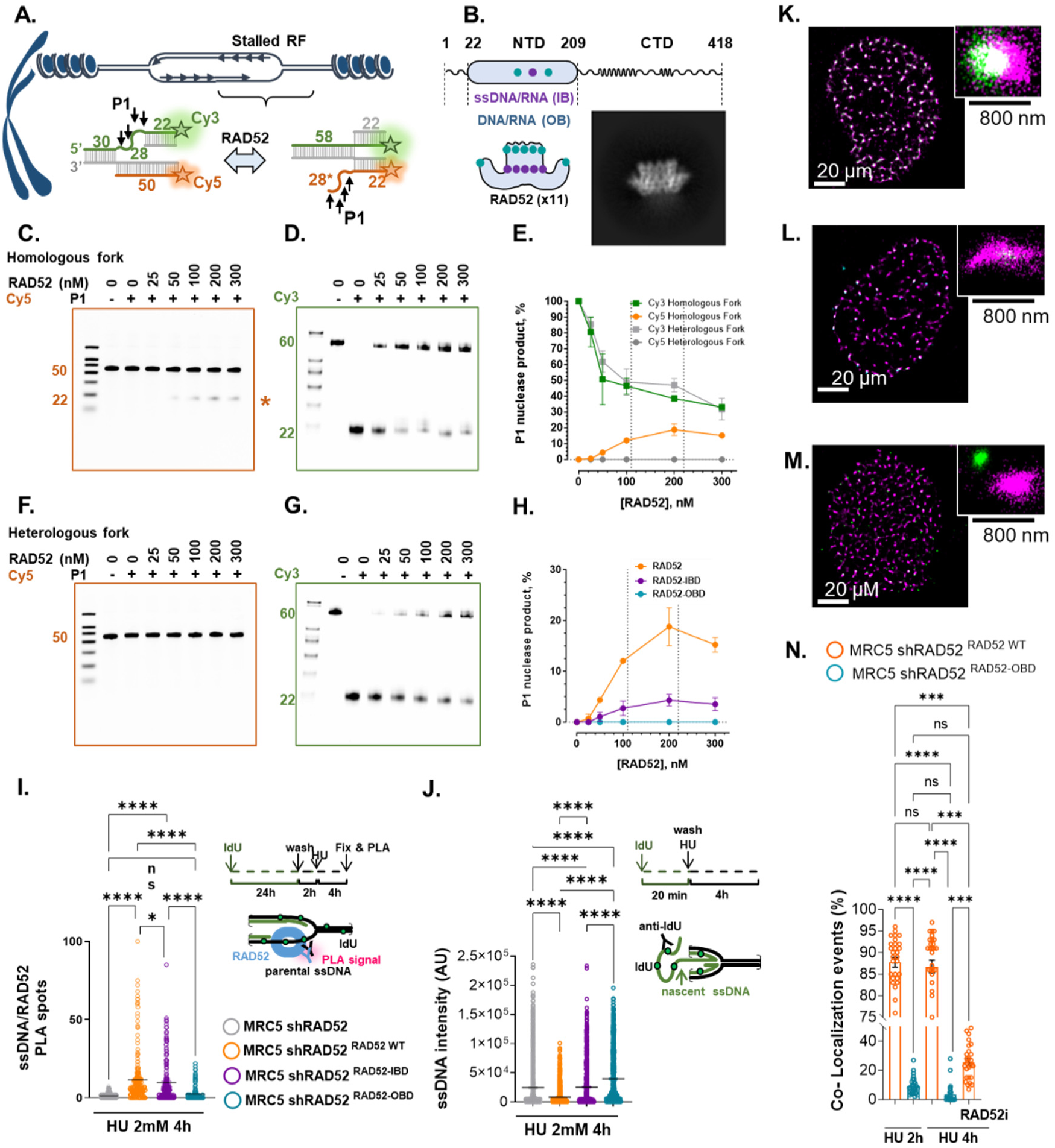
RAD52 mediates DNA strand exchange reaction at replication fork. **A.** DNA forks were annealed using oligonucleotides #1, #2, #3 and #4 (homologous fork) or #5 (heterologous fork) listed in **Supplemental Table S1**. The anticipated product is shown on the right. **B.** Domain organization of the RAD52 monomer and the side view of the structured RAD52 undecameric ring formed by its N-terminal domain (NTD). The inset shows a representative cryo-EM 2D class average of the apo ring of RAD52. **C.** A representative denaturing SDS-PAGE gel showing products of the P1 digestion of the Cy5-labeled strand. Asterisk marks the product dependent on the strand exchange confined to the gap region. **D.** The same as **C,** but for the Cy3-labeled strand. **E.** Quantification of the unique Cy5 (orange) and Cy3 (green) bands is shown as average and standard deviation for three independent experiments. **F.** and **G.** are the same as **C** and **D**., but for the heterologous fork. **H.** Quantification of the strand exchange product for the reactions containing wild type RAD52 (orange), RAD52^IBD^ (purple) and RAD52^OBD^. The data are shown as average and standard deviation for three independent experiments. **I.** PLA analysis of RAD52 binding to parental ssDNA. RAD52 knockdown cells, complemented with the indicated RAD52 mutants were treated with 100 μM IdU for 20 hours, released for 2 hours in fresh medium and exposed to HU. The PLA reaction was carried out using antibodies against RAD52 and IdU. The graph reports the number of PLA spot per nucleus. See **Supplemental Figure S2** for representative images and Western blot. ns = not significant; ****P < 0.0001; Kruskal-Wallis test. **J.** Immunofluorescence analysis of the exposure of ssDNA upon replication fork reversal and degradation. RAD52 knockdown cells complemented with the indicated RAD52 version were treated for 4h with HU. The ssDNA was detected by native anti-IdU immunofluorescence. Graph shows the intensity of ssDNA staining (AU) per cell from two replicates (****P < 0.0001; Kruskal-Wallis test). See **Supplemental Figure S2** for representative images**. L. – N.** Representative images from nuclei treated with 4h HU and showing clusters of EdU and EdU-RAD52 detected by dSTORM found in cells expressing wild-type RAD52 (L), RAD52-OBD (M) or treated with EGC (N). The mixed colors denote binary signals. **O**. Quantification of EdU-RAD52 localization by dSTORM. The graph shows % of EdU-RAD52 co-localization events from 20 images. All the values above are presented as means ± SD. (****P < 0.0001; *** p < 0.001 Mann-Witney test).

Each monomer of RAD52 contains two DNA binding sites, so-called primary or inner binding (IB) site located in a groove that spans the RAD52 ring circumference and can only accommodate nucleic acids in the single stranded form, and the secondary or outer binding (OB) site^25,30–32^ (**Fig. 1B**.). In the P1 nuclease assays, the RAD52^IBD^ mutant (K152A/R153A) with the disabled inner DNA binding site displayed a four-fold reduced capacity to sensitize the Cy5 strand to P1 (**Fig.1H**), while the outer binding defective RAD52^OBD^ (K102A/K133A/K169A/R173A) was completely devoid of the strand exchange activity on the fork (**Fig.1H**). The effect of the IBD and OBD mutations on the RAD52-mediated strand exchanged was paralled by their effect on the RAD52-mediated protection of replication forks from both reversal and restoration by SMARCAL1 *in vitro* with outer binding site being more important for both activities than the inner binding site (**Supplemental Fig. S2&3**).

In cells, RAD52 binds stalled DNA replication forks protecting them from excessive reversal by SMARCAL1, followed by MRE11-dependent degradation and exposure of nascent DNA in the detectable single-stranded form^4,5^. To investigate if altered DNA strand exchange activity and fork protection detected *in vitro* also results in defective fork binding and protection in cells, we added-back the wild-type or mutant RAD52s to cells stably-expressing an shRAD52 cassette directed against the 3’ UTR^5^. As adding-back RAD52 leads to elevated protein expression, we equalized the expression of the wild type and mutant RAD52 by combining transfection with an shRAD52-resistant RAD52 plasmid and a RAD52 siRNA targeting the CDS^18^ (**Supplemental Fig. S4A**). We first analyzed the association of the two RAD52 mutants with parental ssDNA at stalled replication forks using proximity ligation (PLA). **Fig.1I** and **Supplemental Fig. S4B&D** show that RAD52 is recruited to the parental ssDNA in hydroxyurea (HU) treated cells, confirming our previous observations^5,25^. In contrast, both the RAD52^IBD^ and the RAD52^OBD^ mutants showed defects in association with parental ssDNA, the RAD52^OBD^ being more severely affected. As expected, RAD52-depleted cells showed increased levels of nascent ssDNA upon replication fork arrest^5,25^. Adding-back wild type RAD52 restored the normal levels of exposed ssDNA. RAD52^IBD^ only partially complemented the RAD52 depletion, while RAD52^OBD^ was completely unable to revert the exposure of nascent ssDNA seen in the absence of RAD52 (**Fig.1J** and **Supplemental Fig. S4C&E**).

To provide further evidence of the outer DNA binding site importance to RAD52 recruitment to the perturbed forks in response to replication stress, we performed super-resolution microscopy by dSTORM. Nascent DNA at the fork was labeled by a short EdU pulse and ectopically expressed RAD52 (wild type or RAD52^OBD^) was detected via the appended HA tag (**Fig. 1K-N**). Cells were subjected to dSTORM and co-localization clusters were evaluated after filtering out background and artefactual co-localization arising from cross-excitation of the labels. As shown in **Fig. 1K-M**, RAD52 was easily found clustering with EdU at nanoscale and co-localization events increased with time in HU and, notably, were reduced by treatment with the RAD52 inhibitor EGC^22^. RAD52 clustering at EdU-labelled replication forks was dramatically reduced in the RAD52^OBD^ mutant. Inspection of the nanoscale topology of binary co-localization at 50-80 nm showed proximity of the EdU-stained DNA with the RAD52 signal suggesting close association at the end of the labelled site, *i.e.,* near the fork. This nanoscale spatial organization was lost upon treatment with EGC.

### DNA replication fork enforces a unique two-ring RAD52 architecture

To quantify the stoichiometry of the RAD52-fork interaction we employed mass photometry, a light scattering based single-molecule technique^41,42^. In the absence of DNA, RAD52 exists in solution as a mixture of undecamers, decamers and monomers^25^ (**Fig.2A**, **Supplemental Table S2**). The undecameric rings of RAD52 are readily observed in the 2D classes of the cryo-EM analysis described below (**Fig. 2B**). Addition of the replication fork like DNA structure (homologous or heterologous fork with a 30-nucleotide leading strand gap) results in the new peaks corresponding to RAD52-fork complexes. Increasing concentration of RAD52 populates peaks corresponding to two and to a lesser degree three undecamers of RAD52 per fork (**Fig.2C&D**, **Supplemental Table S2**). Double rings bound to the fork DNA were also observed in the RAD52^IBD^ mutant, but were much reduced in the RAD52^OBD^ mutant despite its capacity to bind DNA^25^ (**Supplemental Fig. S5**, **Supplemental Table S2**). RAD52 was also able to bind to and to form double-ring structures on the fork DNA bound by RPA (**Supplemental Fig. S6**, **Supplemental Table S3**).

**Figure 2.**
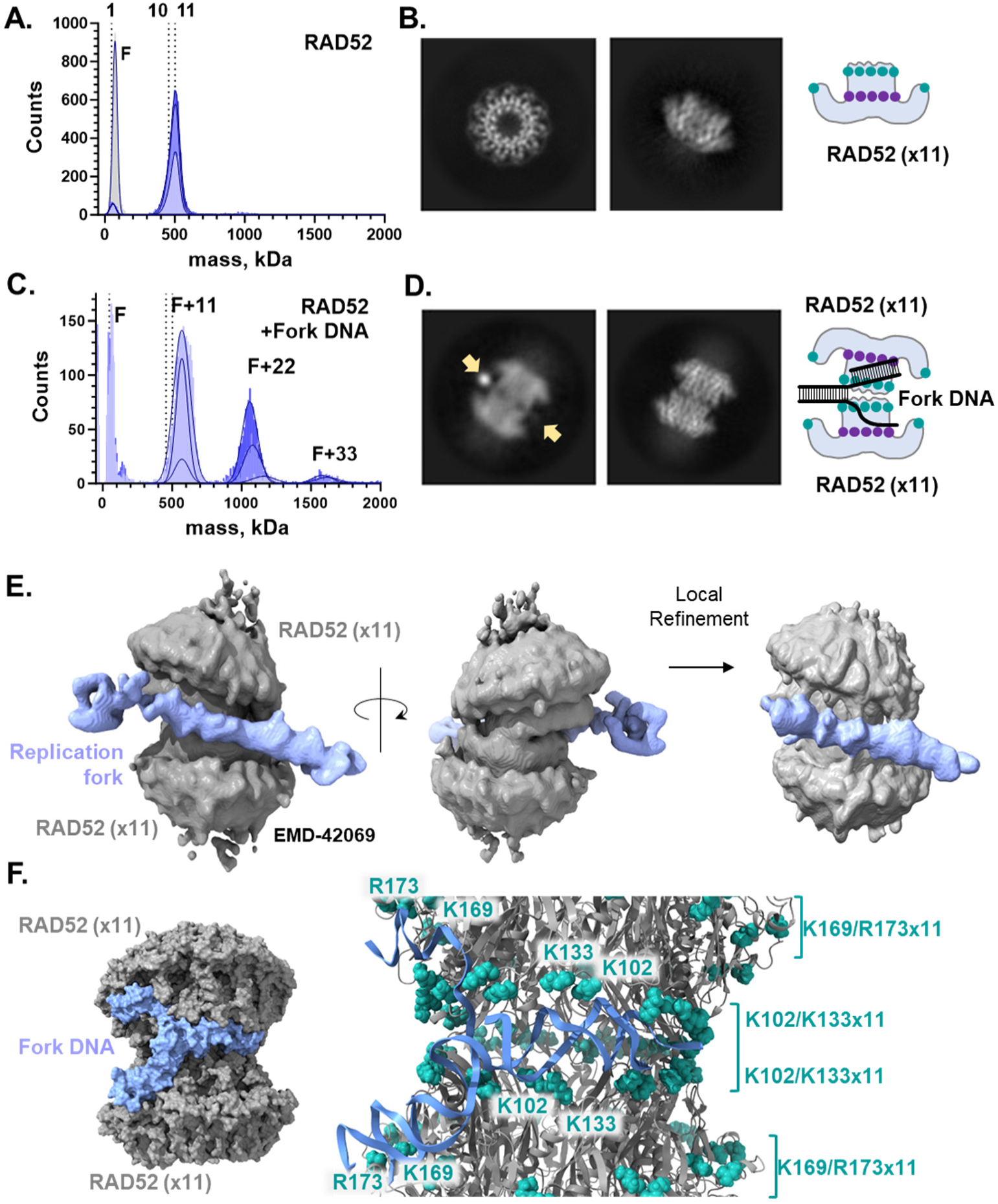
RAD52 is organized on the DNA replication fork into a unique two-ring structure. For all experiments, the heterologous fork was assembled from oligos #2, #3, #4 and #6 listed in **Supplemental Table S1**. **A.** Mass photometry experiment showing oligomeric states of 110, 220 and 330 nM RAD52 (from light to dark blue) and DNA fork (grey) (see **Supplemental Table S2** for quantification). **B.** Representative 2D classes from the cryo-EM analysis of the apo RAD52 ring depicting the top and side views. The cartoon representation shows the location of the inner (purple) and outer (teal) DNA binding sites (see **Supplemental Figure S7** and **Supplemental Table S4** for the 2.5 Å resolution structure and cryo-EM data collection and processing workflow). **C.** Mass photometry experiment showing oligomeric states of 110, 220 and 330 nM RAD52 (from light to dark blue) bound to the fork DNA (see **Supplemental Table S3** for quantification). **D.** Representative 2D classes from the cryo-EM analysis of the RAD52-fork complex showing a side view with the DNA visible (marked by arrows) and another side view with two rings slightly off register. See **Supplemental Figure S8-9** for the structures and cryo-EM workflow. **E.** A cryo-EM structure of the RAD52-fork DNA complex. The RAD52 double undecamer is shown in grey and the fork DNA is in blue (see **Supplemental Figure S8-9** and **Supplemental Table S4** for the additional structures, and cryo-EM data collection and processing workflow). **F.** All atom model showing placement of a smaller model fork into the RAD52 double ring (see Methods section for details, and **Supplemental Figure S11**). In the zoom-in, the residues in the OB sites are shown in teal and the DNA is shown as blue ribbons.

To understand the nature of the RAD52 double-undecamer induced by the replication fork, we analyzed it by cryo-EM (**Fig. 2**; **Supplemental Table S4**; **Supplemental Figures S7-S9**). Similarly to the published crystal structures of the RAD52 N-terminal domain^32,35,36^ and cryo-EM structures of the full-length human RAD52^37,38^, our 2.5Å structure of the RAD52 in apo form also showed an undecameric RAD52 N-terminal domain ring (RMSD value of 0.49 Å between apo form and PDB code of 1KN0), and a disordered C-terminus (**Supplemental Fig. S7**). In the presence of the replication fork DNA, RAD52 assembled in an unexpected and unique two-ring structure (**Fig.2D&E**, **Supplemental Figure S8-9**). In it, the two rings of RAD52 are facing head-to-head, either slightly shifted or fully aligned, forming a spool-like arrangement (**Supplemental Figure S8**).

The residues comprising the outer binding site are arranged along the spool’s central shaft forming a continuous surface that can accommodate the two arms of the fork undergoing strand exchange. DNA duplex, likely the parental strand, enters the shaft from the side. While mass photometry experiments multiple rings of RAD52 can also bind to a long ssDNA or dsDNA, the spool-like organization is rarely observed on these substrates in single particle cryo-EM analyses (**Supplemental Figure S10**).

To understand how the fork organizes the RAD52 rings into the double-ring spool poised for strand exchange, we constructed an all-atom model of the RAD52 double-ring system by fitting the two high resolution undecameric rings of RAD52 derived from our apo structure into the low resolution density map for the RAD52-replicaiton fork complex. The model was optimized using a refinement method that employs the YASARA knowledge-based force field and a simulated annealing MD protocol (see Methods; **Supplemental Figure S11**). A small fork DNA was built in MOE based on the structure of the four-way junction (PDB: 1XNS) and optimized using an accelerated MD sampling approach LowModeMD^43^. The lowest energy version of the fork structure was employed in a rigid body docking into the RAD52 two-ring model using AutoDock VINA. The best docking pose was then subjected to another round of the LowModeMD conformational search (see Methods; **Supplemental Figure S11**). The model fork engages both rings of RAD52 making extensive interactions with the key residues of the bipartite outer binding site (**Fig.2F**) highlighting the importance of this site to the two-ring structure formation and strand exchange at the fork and explaining the involvement of the K169 and R173 residues. While no contacts were observed with the inner binding site, our small model fork used in the computational analysis contained short arms and represented the initial formation of the two-ring RAD52 arrangement. A longer ssDNA (gap or displaced strand) is expected to engage with the inner binding site as well.

### The nature of the stalled fork influences the dynamics of the strand exchange reaction

To evaluate the directionality and dynamics of the RAD52-mediated strand exchange at the replication fork we utilized single-molecule FRET (smFRET) enabled by the total internal reflection fluorescence microscopy (TIRFM). While RAD52 can facilitate DNA strand exchange^33,44^, this reaction is distinct from the eponymous reaction carried out by the RecA-family recombinases, whose directionality depends on the energy of ATP binding and hydrolysis. To observe the fork rearrangement by RAD52 we constructed a homologous fork with the 30 nucleotide leading strand gap. The Cy5 dye (FRET acceptor) was placed at the 3’ end of the nascent leading strand and the Cy3 dye (FRET donor) was placed opposite Cy5 on the lagging strand gap (**Fig 3A**). The fork was tethered to the TIRFM flow chamber via biotin moiety located at the parental duplex (**Fig 3A**). In the absence of protein, the two dyes produce stable fluorescence and FRET signal around 0.26 with no fluctuations (**Fig 3B&C**). RAD52-mediated strand exchange rearranges the folk bringing Cy3 and Cy5 close to one another resulting in increase in FRET up to 0.81 (determined by constructing the DNA substrate corresponding to the product of the strand exchange reaction, *i.e.* 4-way junction in Fig 3). In the presence of RAD52, the strand exchange product builds over time and the individual trajectories show slow fluctuations indicative of the bidirectional strand exchange (**Fig 3B&C**). While RAD52 binding results in some rearrangement of the heterologous fork incompatible with the strand exchange reaction, the high FRET corresponding to the completed strand exchange is not observed. The presence of RPA on the ssDNA gap decreases the observed FRET but does not interfere with the strand exchange (**Supplemental Fig. S12**).

**Figure 3.**
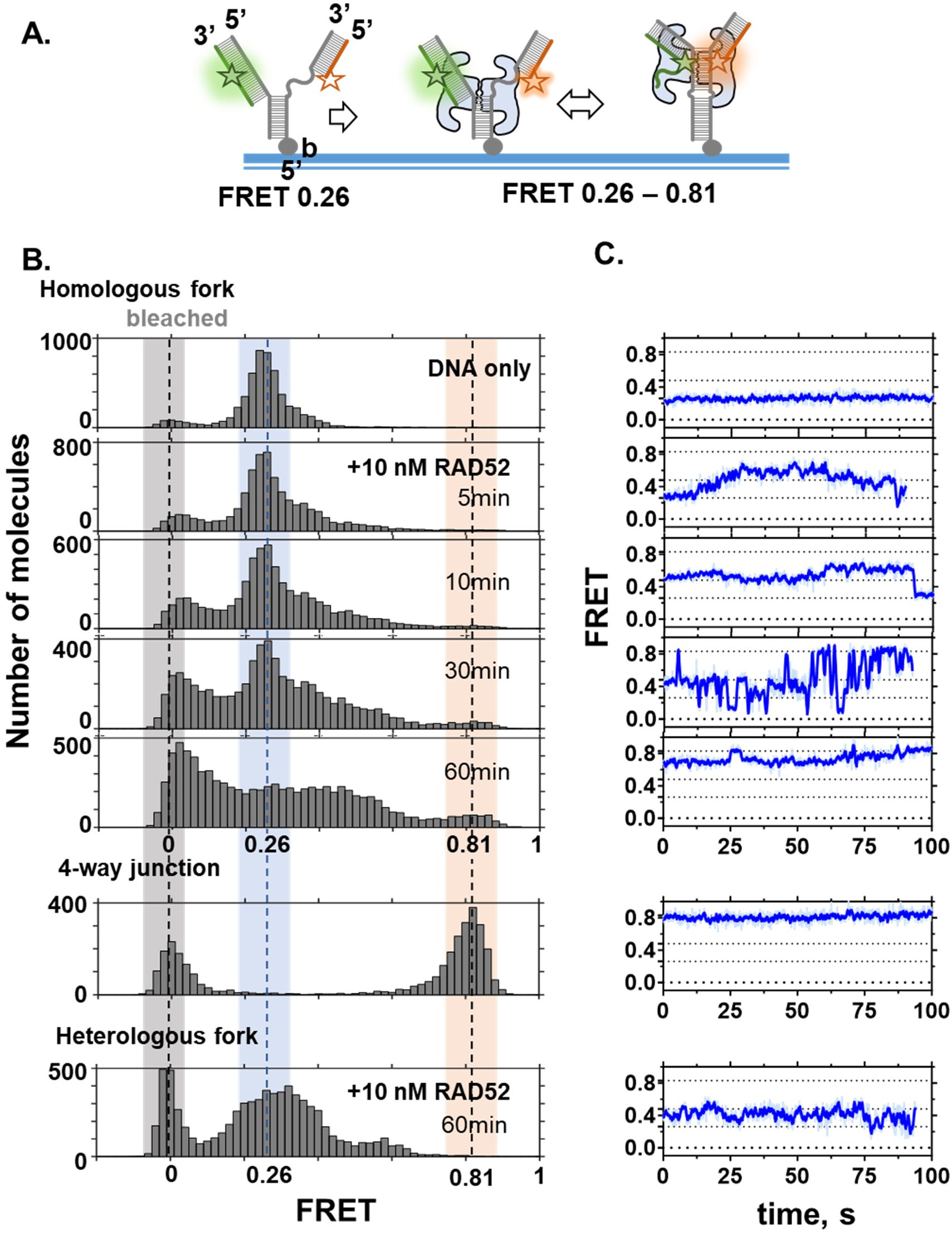
Single-molecule FRET analysis of the RAD52-mediated strand exchange reaction on the model replication fork. **A.** Cartoon depiction of the experimental design. A model replication fork with a leading strand gap was assembled from oligonucleotides #2, #3, #4 and #6 listed in **Supplemental Table S1** and schematically depicted in **Supplemental Figure S1**. The Cy3 (FRET donor) and Cy5 (FRET acceptor) dyes are placed at the lagging and leading arms of the fork (FRET 0.26). Complete strand exchange reaction confined to the gap region yields a 4-way junction and a characteristic FRET 0.81). **B.** Single-molecule FRET distributions for the forks containing two fully homologous arms (homologous fork), fully exchanged fork (4-way junction), and heterologous fork (the ssDNA gap region consists of 30 Ts). Note that the peak around 0 FRET corresponds to the molecules with bleached Cy5 dye which accumulate over time. **C**. Representative smFRET trajectories (time-based changes in the FRET signal from individual fork molecules) for each condition in **B**.

### RAD52 directly competes with SMARCAL1 at the replication fork

To determine the consequences of the RAD52 fork interaction on SMARCAL1 binding we carried out mass photometry experiments in the presence of SMARCAL1 and heterologous fork DNA containing 30 nt gap on the leading strand (**Fig.4A-E**; **Supplemental Table S5**). SMARCAL1 (∼107 kDa) readily formed 1:1 complex with fork DNA (∼75 kDa) (**Fig.4B**). At low RAD52 concentrations corresponding to one undecamer per fork, we observed peaks corresponding to RAD52-fork, SMARCAL1-fork complex and ternary complex of both proteins bound to the fork. Increasing the RAD52 concentration to two or three undecamers per fork resulted in the disappearance of the SMARCAL1-fork peak and appearance of the peak corresponding to two RAD52 rings per fork suggesting that formation of the two-ring structure is key for competing with SMARCAL1 for the fork access (**Fig.4B**; **Supplemental Table S5**). Both RAD52 mutants displayed defect in competing with SMARCAL1 with the defect being more profound in the OBD mutant (**Fig.4D&E**) consistent with the defect in these mutants in antagonizing fork reversal and restoration by SMARCAL1 (**Supplemental Figures S1&2**).

**Figure 4.**
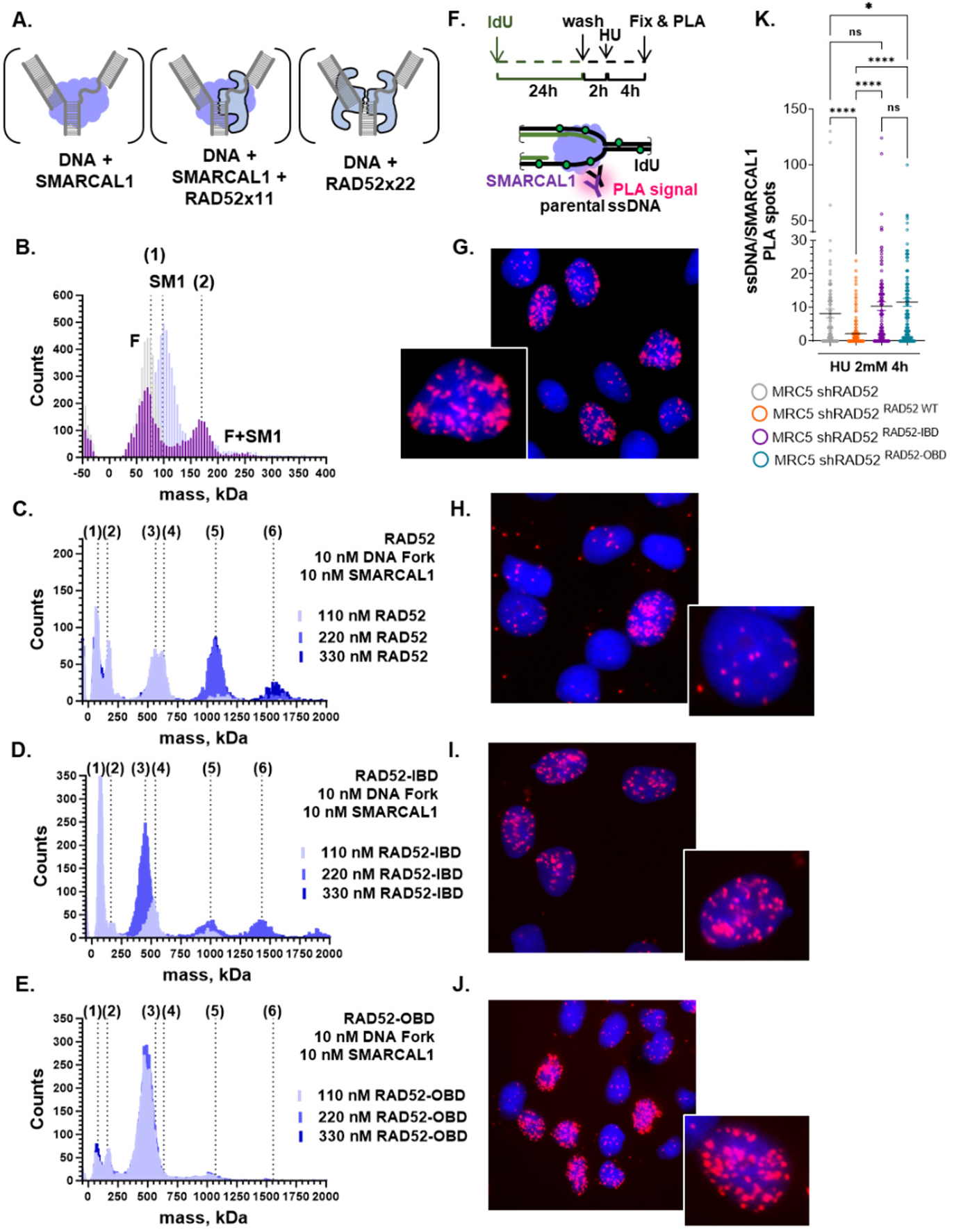
Competition between RAD52 and SMARCAL1 for binding to replication fork depends on the integrity of the outer DNA binding site. **A.** Cartoon depiction of the molecular species expected in the mass photometry (MP) experiment. B. MP analysis of the SMARCAL1-fork interaction. Distributions are shown for fork alone (grey), SMARCAL1 (alone) and the SMARCAL1:fork complex (purple). **B. – E.** Representative MP data for the complexes formed by RAD52, fork and SMARCAL1. (1) Peak corresponding to fork or SMARCAL1 alone; (2) SMARCAL1-fork complex; (3) one ring of RAD52+fork; (4) one ring of RAD52+fork+SMARCAL1; (5) two rings of RAD52+fork; (6) three rings of RAD52+fork. Quantification of the MP data is summarized in the **Supplemental Table S5**. **F.** Schematic of the fork:SMARCAL1 experiment. **G. – J.** Analysis of SMARCAL1-parental ssDNA interaction by PLA. RAD52 knockdown cells, complemented with the indicated RAD52 mutants were treated with 100 μM IdU for 20 hours, released for 2 hours in fresh medium and exposed to HU. The PLA reaction was carried out using antibodies against the SMARCAL1 protein and IdU. Representative images are shown. Magnification of one nucleus is presented in the inset. **K.** Quantification of the number of PLA spot per nucleus. (ns = not significant; **P<0.1; ***P < 0.001; ****P < 0.0001; Kruskal-Wallis test).

Mass photometry suggests competition between RAD52 and SMARCAL1 for fork binding. To confirm this potential competition in cells, we used parental ssDNA PLA to evaluate binding of SMARCAL1 to the parental gap formed at the fork upon HU-induced replication arrest in cells expressing the wild-type RAD52, RAD52^IBD^ and RAD52^OBD^. Depletion of RAD52 resulted in a higher detection of SMARCAL1 at parental ssDNA compared to cells complemented with the wild-type RAD52 (**Fig.4F-K**). Both RAD52 mutants showed increased localization of SMARCAL1 at parental ssDNA (**Fig.4F-K**). Increased SMARCAL1 binding to parental ssDNA stimulated by loss of RAD52 or in cells complemented with RAD52^IBD^ and RAD52^OBD^ was consistently detected at multiple time-points after replication arrest, from 1h to 4h (**Supplemental Fig. S14A, C & D**) and upon delayed replication progression induced by 0.5mM of HU (**Supplemental Fig. S14B**). RAD52 is crucial for viability of BRCA2-deficient cells^3,6,7,11,19–26^. Thus, we investigated if SMARCAL1 localization was increased at parental ssDNA after replication fork arrest when the BRCA2-mutated cancer cells PEO1 were challenged with RAD52 inhibitor EGC during a short HU treatment. SMARCAL1 localization at parental ssDNA was compared with that observed in the BRCA2-revertant clone PEO4. As shown in **Supplemental Fig. S15**, SMARCAL1 is underrecruited in PEO1 compared to PEO4 cells. However, in both the cells, EGC significantly stimulated recruitment of SMARCAL1 at parental ssDNA.

Finally, we performed triple SMLM by dSTORM in U2OS cells labelled with nascent DNA (EdU), RAD52 and SMARCAL1 antibodies to unveil additional signs of competition for the binding to stalled replication forks. Cluster analysis of HU-treated cells confirmed co-localization of RAD52 with fork DNA at the nanoscale level also using the antibody against the endogenous RAD52 and revealed that a consistent number of RAD52 can be detected in triple clusters showing concomitant staining with SMARCAL1 (**Supplemental Fig. S16A&B**). Of note, the proportion of clusters showing EdU-SMARCAL1 increased upon RAD52 inhibition. Visual inspection of the triple-positive clusters revealed that most often RAD52 can be found in the middle of the EdU (DNA) and SMARCAL1 staining, suggesting a competition as evidenced in the *in vitro* experiments.

## Discussion

While a validated and an attractive anticancer drug target (reviewed in ^3^), the RAD52-DNA interaction is complex and agile, complicating selection of the structural features that can be targeted. The DNA binding areas within each RAD52 oligomeric ring are divided into an inner (primary) and outer (secondary) binding sites^25,30–32^. The inner site is a deep narrow groove with positively charged residues making electrostatic contacts with the DNA backbone. Due to its shape, the inner binding site can only accommodate nucleic acids in their single-stranded form^32,35,39^. In the double-ring spool-like structure promoted by binding of RAD52 to stalled DNA replication fork, the two outer binding sites create a continuous surface that connects the two inner binding sites (**Fig. 2; Supplemental Figure S17A**). The two homologous arms can be accommodated on this surface with the ssDNA venturing into the inner binding site. Our Low Mode MD simulation of the initial encounter between the model fork and a two-ring RAD52 structure (**Fig. 2F**) shows the fork interacting primarily with the bipartite outer binding site. Notably, this model explains how the residues on the outer leaflet of the RAD52 monomers (K169 and R173) are involved in the DNA binding. In a larger fork the ssDNA is likely to extend into the inner binding site as well. We envision that the parental ssDNA gap is placed initially in the inner binding site of one of the rings, while the homologous duplex is placed on the barrel-like surface formed by both rings **Supplemental Fig. S17A**. Single-stranded product of the strand exchange reaction is then moved to the inner binding site of the second ring. The reaction can proceed bidirectionally but is limited to the gap region. This constraint is likely enforced by the structure of the fork, as RAD52 has been shown to readily diffuse along both ssDNA and dsDNA while searching for homology during single strand annealing^45,46^. Alternatively, the fork may be splitting at the barrel of the RAD52 double-ring spool promoting fork rearrangement to and from the four-way junction (**Supplemental Fig. S17A**). While the ability of RAD52 to carry out a strand exchange reaction on the unconnected dsDNA and ssDNA^33^ argues for the former scenario, such an arrangement would be inconsistent with a directional placement of ssDNA into the inner binding site adding weight to the second scenario. In both cases, the nucleoprotein arrangement during fork protection (**Supplemental Fig. S17A**) is different from that proposed for the RAD52-mediated single-strand annealing reaction based on the structural and biochemical data^32,39,45^ (**Supplemental Fig. S17B**).

Collectively, our structural and dynamics data highlight an extraordinary complexity and plasticity of the RAD52-DNA interaction depicting both structural and functional differences of the RAD52-fork complexation and rearrangement from other activities of this multifunctional protein, with the two-ring spool-like arrangement of the RAD52 providing new insights that were not obvious from the previous structures. We propose that binding to the stalled or damage replication fork and its rearrangement through the RAD52-mediated strand exchange converts the fork into a nucleoprotein complex refractory to the fork reversal motors, such as SMARCAL1. RAD52 dissociation would allow branch migration of the fork back to the original state accessible to SMARCAL1. A persistent RAD52-fork complex may eventually recruit MUS81 nuclease to introduce a double-strand break^18^ and initiate fork repair by homologous recombination.

There are several similarities and differences between the RAD52-mediated strand exchange at the replication fork and the eponymous reaction mediated by the RecA-family recombinases, such as RAD51. The most significant difference is that the RAD52-mediated strand exchange is confined to the gap region and is not followed up by the branch migration. Recently, the ability to carry out DNA and DNA/RNA strand exchange reaction was found in the intrinsically disordered DNA binding region of tumor suppressor PALB2 (partner and localizer of BRCA2) extending the range of facilitators of this important cellular process^47,48^. Here we show that the reaction can also occur on the surface of the unique spool-like structure formed by the two undecameric rings of the DNA repair protein RAD52. Akin PALB2-mediated strand exchange, this reaction is different in its mechanism from that catalyzed by the canonical RecA-family recombinases. In contrast to both, however, the RAD52-mediated strand exchange readily proceeds in the presence of RPA.

## Methods

### Chemicals and reagents

All chemicals were reagent grade (Sigma-Aldrich, St. Louis, MO). Cy3-labeled, Cy-5 labeled, biotinylated and unmodified oligonucleotides were purchased from Integrated DNA Technologies. Sequences of all DNA oligonucleotides are listed in **Supplementary Table S1**. Twelve mM Trolox (6-hydroxy-2,5,7,8-tetramethylchromane-2-carboxylic acid, Sigma-Aldrich; 238813-1G) solution was prepared as described previously by adding 60 mg of Trolox powder (238813-5G, Sigma-Aldrich) to 10 mL of water with 60 μL of 2 M NaOH, mixing for 3 days, filtering, and storing at 4 °C. Oxygen scavenging system, Gloxy was prepared as a mixture of 4 mg/mL catalase (C40-500MG, Sigma-Aldrich) and 100 mg/mL glucose oxidase (G2133-50KU, Sigma-Aldrich) in K50 buffer.

### Proteins

The 6xHis-tagged human RAD52 protein (and mutants), Flag-tagged SMARCAL1 and untagged human RPA were expressed and purified using established protocols^5,22,25,28,49^ with a minor modification: SMARCAL1 was concentrated, and buffer exchanged 5 times to storage buffer containing 50 mM HEPES (pH 7.5), 100 mM NaCl, 0.5 mM EDTA, 10% glycerol and 1 mM DTT using Amicon Ultra-0.5 ml (10 kDa MWCO, Millipore) to remove x3 FLAG peptides. Concentrated SMARCAL1 was immediately divided into small aliquots and preserved at −80°C. Protein concentrations were determined using extinction coefficients of 40,470 M^-1^cm^-1^ (RAD52), 88,830 M^-1^cm^-1^ (RPA) and 98,700 M^-1^cm^-1^ (SMARCAL1), respectively.

### DNA substrates

The PAGE-purified Cy3, Cy5 and biotin-labeled oligonucleotides (see **Supplemental Table 1**) were custom synthesized by IDT. DNA substrates mimicking stalled or reversed DNA replication forks were prepared by annealing. The design of the fork for each experiment is schematically illustrated in Supplemental Figure S1. The four-way junction used in the fork restoration assays is based on the ^50^. The leading or lagging parental strands were annealed with their corresponding nascent strands by mixing the respective oligonucleotides together at the final concentration of 1 μM each in buffer containing 10 mM Tris-HCL (pH 8.0), 1 mM EDTA and 150mM NaCl, heated to 95°C for 10 min in a thermocycler and then slowly cooled to 60°C. Then, equal amounts of leading and lagging DNA structures were mixed and slowly cooled to 4°C to complete annealing of the parental-parental region. Annealed forks were used without further purification in cryo-EM studies. For all other analyses, fully annealed fork DNA substrates were separated by PAGE and purified using Model 442 electro-eluter system (BIO-RAD) and stored at 4°C until use.

### P1 nuclease mapping

10 nM of purified fork DNA was mixed with the indicated concentrations of RAD52 (wild type or mutants) in 10 μl of standard reaction buffer, containing 20 mM Tris-Acetate (pH 7.5), 150 mM NaCl, 5 mM magnesium acetate, 1 mM DTT and 0.1 μg/ml BSA and incubated for 30 min at 25°C. Five units of P1 nuclease (NEB) were added, and the reactions were further incubated for 30 min at 25°C. Reactions were stopped and deproteinized by the addition of 10 μl of solution containing 0.5% (v/v) SDS and 0.5 mg/ml proteinase K and incubation at room temperature for 60 min. The 20 μl reactions were quenched by the addition of 15 μl of stop solution containing 10 M Urea, 20% glycerol, 0.1% Orange G. The reaction products were separated by electrophoresis on a 10% (19:1) native polyacrylamide gel in 1× TAE buffer containing 40 mM Tris-Acetate (pH 8.0) and 1 mM EDTA for 2 h at 50 V at 50°C. The gel was imaged using a ChemiDoc (Bio-Rad) by exciting and monitoring Cy3 and Cy5 fluorescence.

### FRET-based fork reversal and fork restoration analysis

FRET-based analyses of DNA fork regression by SMARCAL1 in the presence of RPA and RAD52 (wild type or mutants) were carried out using a Cary Eclipse fluorescence spectrophotometer (Agilent technologies) at 30°C in buffer containing 20 mM HEPES (pH 7.5), 100 mM NaCl, 5 mM MgCl_2_, 100 μg/ml BSA, 2 mM ATP and 1 mM DTT as described previously^5,51^. Briefly, measurements began with buffer only (baseline) followed by addition of the 5 nM of the homologous fork DNA substrate dually labeled with Cy3 and Cy5 fluorophores. The Cy3 fluorophore was excited at 530 nm, and the emission of the acceptor Cy5 and donor Cy3 fluorophores were monitored simultaneously at 660 nm and 565 nm, respectively. After pre-incubation with 10 nM RPA and/or 165 nM RAD52 for 5min, fork regression reaction was initiated upon by addition of the 0.5 nM SMARCAL1. The FRET signal was calculated from the Cy3 and Cy5 fluorescence at each data point as *FRET* = (*I*_Cys_ ∗ 4.2)/(*I*_Cy3_ ∗ 1.7 + *I*_Cys_ ∗ 4.2), where *I*_Cy3_ and *I*_Cys_, are background corrected intensities of the two dyes. For each experiment, the FRET vs. time progress curves were plotted using GraphPad Prism 7.0 and fitted to single exponential decay equations. The rates of the fork reversal in units of nM DNA per minute per nM SMARCAL1 were calculated as *v* = (*k* ∗ *span*)⁄*span_SMARCAL1_*〉 ∗ 3 *nM* ∗ 60, where *k* is the exponential decay constant, *span* is the FRET change between the substrate and the products, and 〈*span*_SMARCAL1_〉 is the average span for the reaction containing SMARCAL1 only.

### Single-Molecule TIRFM

Fifty pM of indicated biotinylated, Cy3 and Cy5-labeled DNA substrates in T50 buffer (10 mM Tris (pH 8.0), 50 mM NaCl) were immobilized on the surface of the microscope quartz slides (Fisher Scientific), which were pre-coated with PEG and bio-PEG to eliminate nonspecific surface adsorption of proteins. The immobilization was mediated by biotin-neutravidin interaction between biotinylated DNA, neutravidin (Thermo Fisher), and biotinylated polymer (PEG-MW 5,000, Nectar Therapeutics). The standard buffer contained 10 mM Tris-Acetate (pH 7.5), 150 mM NaCl, 5mM MgCl2,1 mM DTT and the oxygen scavenging system consisting of 1 mg/ml glucose oxidase (Sigma), 0.4% (w/v) D-glucose (Sigma), 0.04 mg/ml catalase (EMD biosciences) and 1 mM 6-hydroxy-2,5,7,8-tetramethyl-chromane-2-carboxylic acid (Trolox) (Sigma Aldrich). After DNA was tethered on the surface, the indicated amount of RAD52 and/or RPA were added and incubated for indicated amounts of time at 25°C in the standard buffer before starting the recording. Custom built, prism type TIRFM was used to excite fluorophores present on the DNA molecules. Cy3 fluorophores were excited by a DPSS laser (532 nm, 75mW, Coherent), while the Cy5 fluorophores were excited via FRET from Cy3. The fluorescence signals originated from the Cy3 and Cy5 dyes were collected by a water immersion 60× objective (Olympus), separated by a 630 nm dichroic mirror, passed through a Cy3/Cy5 dual band-notch filter (Semrock, FF01-577/690) in the emission optical path. Images were further filtered by using a Chroma ET605/70m filter (for Cy3 emission) and a Chroma ET700/75m filter (for Cy5 emission) inside the dual-view system (DV2; Photometrics) and detected by the EMCCD camera (Andor) with a time resolution of 100 ms. Single-molecule trajectories were extracted from the recorded video file by a custom written IDL code (generously shared by the Taekjip Ha lab, Harvard). Fluorescence trajectories were analyzed using customized MATLAB (The MathWorks, Inc.) scripts (available upon request from the Spies lab). To generate the FRET efficiency histograms, ten movies with a duration of approximately 20 seconds were recorded in different regions of the slide chamber. FRET values were collected from at least 5,000 molecules for each condition as previously described^52^. FRET efficiency histograms were plotted using GraphPad Prism software and fit to multiple Gaussian peaks. A zero FRET peak (3∼30% of total population) represents molecules with the photo-bleached FRET acceptor (Cy5).

### Mass Photometry (MP)

All MP experiments were carried out using the Refeyn TwoMP mass photometry instrument (Refeyn Ltd. Oxford, UK). Molecular weight calibrations were performed using solutions of β-amylase (56, 112 and 224 kDa) and Thyroglobulin (670 kDa). In each experiment, fork DNA, RAD52, SMARCAL1 and/or RPA were sequentially added into buffer-filled gasket containing 30 mM Tris-Acetate pH 7.5, 150 mM NaCl, 5 mM MgCl_2_ and 1 mM DTT to achieve final concentrations as indicated. In the reaction containing cross-linked RAD52-fork DNA species, the complexes were prepared at 5.5 µM and 2 µM concentration of RAD52 (monomers) and fork (molecules), respectively, incubated with 2 mM BS3 for 15 minutes on the ice, and diluted 20 times into buffer-filled gasket (30 mM Tris pH 7.5, 75 mM KCl, 5 mM MgCl_2_ and 1 mM DTT). All experiments were carried out at room temperature. Individual molecular weights collected in a 3000-frame movies (59.9 seconds) were binned in 5 kDa bins and plotted as frequency histograms. GraphPad Prism and DiscoverMP software (Refyen) were used to fit the molecular weight distributions to multiple Gaussians.

### Cryo-EM Sample and Grid Preparation

Cryo-EM sample and grid preparation of the apo RAD52 protein diluted in a buffer containing 20 mM Tris pH 9.0, 200 mM KCl, 1 mM DTT to the final concentration of 0.25 mg/mL (5.2 μM) and stored at 4 °C until grid preparation. Cryo-EM grid for apo RAD52 was generated by applying 3.5 μL of sample to a Quantifoil R0.6/1 300 mesh copper grid for tilted data collection and Ultra AU Foil R1.2/1.3 300 mesh gold grid for un-tilted data collection. The grids were glow-discharged using a PELCO easyGlow for 60 seconds. The grids were blotted for 3 seconds at 4 °C and 100% humidity before plunge-frozen in liquid ethane using an FEI Vitrobot Mark IV.

Cryo-EM and grid preparation of the RAD52-Fork DNA complex prepared in two grids. The complex diluted in a buffer containing 20 mM Tris pH 7.5, 75 mM KCl, 5 mM MgCl_2_, 1 mM DTT. The samples prepared by mixing RAD52 (5.5 µM) and Fork DNA (2 µM) and storing on ice for 20 minutes, followed by addition of BS3 to the final concentration of 2 mM, then mixture is stored on ice for 15 minutes. The first cryo-EM grid was generated by applying 3 µL of sample (2-fold diluted) to a Quantifoil R1.2/1.3 300 mesh Au grid. The grid was hydrophilized / plasma cleaned with a mixture of H_2_ and O_2_ gas (6.4:27.5 ratio) at 50 W for 30 seconds in a Solarus Model 950 Advanced Plasma System (Gatan). The grid was blotted for 4 seconds at 4 °C and 100% humidity before plunge-frozen in liquid ethane using an FEI Vitrobot Mark IV. The second cryo-EM grid was prepared by applying 3.3 µL of the sample (no dilution) to an Ultra AU Foil R1.2/1.3 300 mesh Au grid. The grid was glow-discharged using a PELCO easyGlow for 60 seconds. The grid was blotted for 3 seconds at 4 °C and 100% humidity before plunge-frozen in liquid ethane using an FEI Vitrobot Mark IV.

### Cryo-EM data collection, processing and model building

Cryo-EM data collection of apo RAD52 was performed at the Pacific Northwest Center for Cryo-EM (PNCC) using serialEM. The data set for apo RAD52 was collected using an FEI Titan Krios 300 kV cryo-electron microscope equipped with a BioContinuum K3 direct electron detector. The data set for RAD52-fork DNA complex was collected using a Titan Krios equipped with TFS Falcon IV direct electron detector at National Center for Cryo-EM Access and Training (NCCAT). For the apo RAD52, two datasets were collected over ∼48 hours from two separate cryo-EM grids. 5602 movies (no tilt) and 5688 movies (30 deg tilt) were combined and processed together. The data were collected using super-resolution mode with a pixel size of 0.4128 Å, a defocus range of –0.8 µm to –2.2 µm, and a total electron dose of 50 e^-^/Å^2^. For the RAD52-fork DNA dataset, a total of 6093 movies were recorded over ∼24 hours from 2 cryo-EM grids. The data were collected with a pixel size of 0.959 Å, a defocus range of –0.8 µm to –2.5 µm, and a total electron dose of 50 e^-^/Å^2^.

All cryo-EM data processing was carried out using cryoSPARC^53^. The beam-induced drift was corrected using cryoSPARC patch motion correction and contrast transfer function (CTF) was carried out using cryoSPARC patch CTF-estimation. The micrographs were manually curated and were used to perform blob picking to generate templates for automated template picking. For apo RAD52, after template picking, multiple rounds of 2D classifications were carried out to generate final particle stacks. Non-uniform refinement in cryoSPARC v3.3.2 was used to obtain the final reconstruction for apo RAD52 with C11 symmetry. An existing structure of RAD52 (PDB 1KN0) was fit into the map and used for iterative building and refinement in Coot and phenix.real_space_refine^54,55^. Map sharpening was performed using DeepEMhancer^56^. After template picking, multiple rounds of 2D classifications were carried out to generate final particle stacks containing double rings. The 2D classes of double rings were used as a template for template picking, and particle extraction for RAD52-fork DNA structures of EMD-42066 and EMD-42069. Multiple rounds of 2D classifications were performed and 2D classes of double rings selected for final particle stacks. Four *Ab-initio* models were generated followed by heterogeneous refinement (4 classes) and reconstructions were obtained after a final homogeneous refinement of two major classes (EMD-42066 and EMD-42069). The templates were created from two maps and used for particle picking and extraction for EMD-42440 structure. Multiple rounds of 2D classifications and one round of heterogeneous refinement were performed to create final particle stacks containing single- and double-ring structure. Then Non-Uniform refinement and 2D classification were performed to create final particle stacks of MD-42440’s ab-initio and final homogeneous refinement. The global resolutions for all structures were determined using the gold standard Fourier shell correlation (GSFSC) 0.143 cut off and map quality evaluated using PHENIX mtriage^57^. The RAD52-fork DNA particle stacks were subjected to local refinement in cryoSPARC. Masks were generated around one RAD52 ring (designated as a top ring) and the second ring including fork DNA (bottom ring-fork DNA), which was followed by local refinement. The component maps of the local refinement jobs were combined by taking the maximum value at each voxel using UCSF ChimeraX. A portion of the software used was curated by SBGrid^58^.

### Computational Modeling of the three-way DNA junction bound to the two-ring RAD52 structure

A model consisting of a small fork structure was designed based on the Holiday junction structure 1XNS^59^ (**Supplemental Figures S7; Supplemental Table S1 oligos #16-18**). The solution structure of the fork was optimized using an accelerated MD sampling approach LowModeMD^43^, implemented in the Molecular Operating Environment (MOE) release 2020.0901. This method concentrates kinetic energy on low-frequency vibrational modes, to populate conformations in multiple low-energy states with high-computational efficiency, and is particularly appropriate for complex systems with large numbers of non-bonded interactions. The LowModeMD conformational search procedure includes an iterative process of initial energy minimization, filtering of high-frequency vibrational modes, a short MD and saving distinct structures in a database. The energy minimization gradient threshold was 0.001 kcal/mol/Å, and searches were configured to terminate after 500 contiguous failed attempts to generate novel conformations, with up to 10 000 iterations. Conformations were identified as unique if their root-mean-square-distance was above a threshold value of 0.25 Å. The LowMode sampling converged on 4 different fork structures, all containing a “corkscrew” like form (**Supplemental Figures S7D**).

An all-atom model of the RAD52 dual ring system was constructed by placing the two undecameric RAD52 rings from the apo structure (PDB 8TKQ) into the cryo-EM density map of EMD-42069 and applying a refinement method that employs the YASARA knowledge based force field^60,61^ and a simulated annealing MD protocol, which has been described in detail here^62,63^. The model with the highest average Z-factor of 0.06 (Energy = -1240209.38; normality of dihedral bonds = 1.98; 1D Packing = -1.30; 3D Packing = -0.51) was selected.

The lowest energy version of the fork structure and the RAD52 two-ring model were employed in a rigid body docking using AutoDock VINA^64^ implemented in YASARA Structure 22.9.24 (YASARA Biosciences GmbH, Vienna, Austria.). A simulation cell of 100X100X100 Å, and a volume of 1000000 Å^3^ was employed, bisecting approximately one third of the entire RAD52 dual ring system. The LowModMD-optimized fork was subjected to 25 docking runs and ranked by the VINA binding energy scores; clustering structures was performed, such that those differing by at least 5.0 Å heavy atom RMSD after superposing on the RAD52 receptor were considered distinct. This resulted in 18 structures. The top four distinctive docked complexes were all relatively isoenergetic with VINA scores from 23.6 to 21.09 kcal/mol. Complexes 1 and 2 displaced DNA placement with significant portions of the ligand exposed to solvent. Although this may be an intermediate in the overall binding process, complexes 3 and 4 had larger contact surfaces and were consistent with what one expects from the experimental studies of the RAD52 mutants. The RAD52-fork complex 3 was then subjected to LowModeMD protocol as described above for the fork DNA. The RAD52 dual ring remained frozen throughout the LowModeMD conformational search, and a single low-energy peptide conformation was identified.

### In situ PLA assay for ssDNA–protein interaction

The *in situ* PLA (Navinci Diagnostics) was performed according to the manufacturer’s instructions. For parental ssDNA-protein interaction, cells were labelled with 100 μM IdU for 20 hours and then released in the fresh medium for 2 hours. After treatment, cells were permeabilized with 0.5% Triton X-100 for 10 min at 4 °C, fixed with 3% PFA/2% sucrose in PBS 1X for 10 min and then blocked in 3% BSA/PBS for 15 min. After washing with PBS, cells were incubated with the two primary antibodies: mouse monoclonal anti-IdU (Becton Dickinson, 1:50), and rabbit polyclonal anti-RAD52 (Aviva, 1:150). Samples were incubated with secondary antibodies conjugated with PLA probes MINUS and PLUS: the PLA probe anti-mouse PLUS and anti-rabbit MINUS (or NaveniFlex equivalent). The incubation with all antibodies was carried out in a humidified chamber for 1 h at 37 °C. Next, the PLA probes MINUS and PLUS were ligated using two connecting oligonucleotides to produce a template for rolling-cycle amplification. After amplification, the products were hybridized with red fluorescence-labelled oligonucleotide. Samples were mounted in Prolong Gold anti-fade reagent with DAPI (blue). Images were acquired randomly using Eclipse 80i Nikon Fluorescence Microscope, equipped with a Virtual Confocal (ViCo) system. The analysis was carried out by counting the PLA spots for each nucleus, plotted and analyzed using GraphPad Prism.

### Detection of nascent ssDNA by native IdU assay

To detect nascent ssDNA, cells were labelled for 15 minutes with 80 µM IdU (Sigma-Aldrich), then treated with or without HU 2mM for 4 hrs. For immunofluorescence, cells were washed with PBS 1X, permeabilized with 0.5% Triton X-100 for 10 min at 4 °C and fixed in 3% PFA, 2% sucrose in PBS 1X. Fixed cells were then incubated with mouse anti-IdU antibody (Becton Dickinson, 1:80) for 1 h at 37 °C in 1% BSA/PBS, followed by species-specific fluorescein-conjugated secondary antibodies (Alexa Fluor 488 Goat Anti-Mouse IgG (H + L), highly cross-adsorbed—Life Technologies). Slides were analysed with Eclipse 80i Nikon Fluorescence Microscope, equipped with a Virtual Confocal (ViCo) system. For each time point, at least 100 nuclei were analyzed. Quantification was carried out using the ImageJ software and the data were plotted using GraphPad Prism.

### Single-Molecule Localization Microscopy

To detect EdU-RAD52 interaction U2OS cells were transfected with AAVS1-TRE3G-RAD52-3xHA then treated as indicated. For the detection of EdU-RAD52-SMARCAL1, U2OS were treated as indicated. After the treatment, cells were fixed and immunofluorescence was performed as described by Whelan & Rothenberg, 2021. EdU was detected with the Click-iT™ EdU Alexa Fluor™ Imaging Kit (Invitrogen) for 30 minutes at RT. For EdU-RAD52 interaction, EdU detection was performed as above. Coverslips with fixed cells were stored for up to 1 week at 4 °C prior to dSTORM imaging. The fixed cells on coverslips were mounted onto concave slides and dSTORM imaging B3 buffer from Oxford Nanoimaging (ONI, PN #900-00004) was added before imaging. A commercial TIRF microscope (Nanoimager S Mark IIB from ONI, https://oni.bio) with lasers of 405 nm/150 mW, 488 nm/1 W, 561 nm/1 W, and 640 nm/1 W was used to acquire data. Briefly, fluorophore emission was collected by an oil immersion 100x/1.45NA objective and images were acquired at 30 ms exposure time using a Hamamatsu ORCA-Flash4.0 V3 Digital sCMOS camera. The reconstruction of the super-resolution image was conducted by NimOS software from ONI. Localization data were then analyzed using the ONI-CODI platform for drift correction, filtering, clustering, and counting.

### Data availability

Atomic coordinates for the modelled RAD52 apo structure have been deposited with the Protein Data Bank (PDB) under accession number 8TKQ. All cryo-EM maps are available from the Electron Microscopy Data Bank (EMDB) under accession numbers of EMD-41537, EMD-42065, EMD-42066 and EMD-42069. The workflows used for data collection and processing are presented in **Supplemental Figures S5 and S6**. Data, plasmids for protein expression, and code for single-molecule data analysis are available from the corresponding author upon request.

## Author contributions

MH – conceptual and experimental design, proteins, DNA substrates, biochemical, FRET and single-molecule analyses, data analysis and interpretation, manuscript preparation; MR - conceptual and experimental design, proteins, cryo-EM, data analysis and interpretation, manuscript preparation, PG – mass photometry analyses and data interpretation; EM – cell-based analyses, data analysis and interpretation, manuscript editing; G.M - cell-based analyses, data analysis and interpretation; LD – cell-based analyses, data analysis and interpretation, manuscript editing; FAA – super-resolution microscopy, data analysis and interpretation; EAP – biochemical and FRET analyses; AS – biochemical and FRET analyses; BJD – biochemical and FRET analyses; LG – cryo-EM, manuscript editing; NJS – cryo-EM, manuscript editing; MAS – computational analyses and interpretation, manuscript editing; PP – conceptual and experimental design, data interpretation, manuscript editing, funding; MS – conceptual and experimental design, data analysis and interpretation, manuscript preparation, funding.

