## Supplemental Information for "A double-ring of human RAD52 remodels replication forks restricting fork reversal"

‡ Equal lead author contribution

† Contributed equally

§ Present address: Medical College of Wisconsin, Milwaukee, WI

§§ Present address: PAQ Therapeutics, Burlington, MA 01803, USA

### Supplemental Table 1: Oligonucleotides used in this study

| NAME | SEQUENCE | EXPERIMENT |
|---|---|---|
| **#1** (80 nt) | 5'-GGTATATTCAACAGTTCAACGACGAATGTTGGTACTCCAGTGTGTAGCATATTACGAA-CTTTTGAGGCAGGCATGGTAGC-(**Cy3**)-3' | P1 (hom); reversal |
| **#2** (80 nt) | 5'-GCTACCATGCCTGCCTCAAAAGTTCGTAATATGCTACACACTGGAGTACCGGCATTC-GTCGTTGAACTGTTGAATATACC-3' | P1; reversal; MP; cryo-EM |
| **#3*** (50 nt) | 5'-GGTACTCCAGTGTGTAGCATATTACGAACTTTTGAGGCAGGCATGGTAGC-(**Cy5**)-3' | P1; MP; cryo-EM |
| **#4** (22 nt) | 5'-GCTACCATGCCTGCCTCAAAAG-3' | P1; MP; cryo-EM |
| **#5** (80 nt) | 5'-GGTATATTCAACAGTTCAACGACGAATGTTGTGCATAACGTCGCTCCGTAGCTATACG-CTTTTGAGGCAGGCATGGTAGC-(**Cy3**)-3' | P1 (het) |
| **#6*** (80 nt) | 5'-GGTATATTCAACAGTTCAACGACGAATGTTTTTTTTTTTTTTTTTTTTTTTTTTTTTTCTTT-TGAGGCAGGCATGGTAGC-(**Cy3**)-3' | MP; cryo-EM (het) |
| **#7** (80 nt) | 5'-(**biotin**)-GGTATATTCAACAGTTCAACGACGAATGTTGGTACTCCAGTGTGTAGCATATT-ACGAACTTTTGAGGCAGGCATGGTAGC-3' | smFRET |
| **#8** (50 nt) | 5'-GGTACTCCAGTGTGTAGCATATTACGA (i**Cy3**) CTTTTGAGGCAGGCATGGTAGC-3' | smFRET (hom) |
| **#9** (22 nt) | 5'-GCTACCATGCCTGCCTCAAAAG-(**Cy5**)-3' | smFRET |
| **#10** (50 nt) | 5'- TTTTTTTTTTTTTTTTTTTTTTTTTTTT (i**Cy3**) CTTTTGAGGCAGGCATGG-TAGC-3' | smFRET (4-way junction) |
| **#11** (80 nt) | 5'-(**biotin**)-GGTATATTCAACAGTTCAACGACGAATGTTTTTTTTTTTTTTTTTTTTTTTTTTTT-TTTCTTTTGAGGCAGGCATGGTAGC-3' | smFRET (het) |
| **#12** (50 nt) | 5'-GGTACTCCAGTGTGTAGCATATTACGAACTTTTGAGGCAGGCATGGTAGC-3' | reversal (leading strand gap) |
| **#13** (22 nt) | 5'-(**Cy5**)-GCTACCATGCCTGCCTCAAAAG-3' | |
| **#14** (22 nt) | 5'-CTTTTGAGGCAGGCATGGTAGC-3' | reversal (lagging strand gap) |
| **#15** (50 nt) | 5'-(**Cy5**)-GCTACCATGCCTGCCTCAAAAGTTCGTAATATGCTACACACTG-GAGTACC-3' | |
| **#16** (17 nt) | 5'-ATGCTATACGAAGTTAT-3' | computational modeling |
| **#17** (35 nt) | 5'-TATAACTTCGTATAGCATACATTATACGAAGTTAT-3' | |
| **#18** (62 nt) | 5'-(**Cy3**)-TAGGCAATTGCCACGTGTCTATCAGCTGAAGTACAAGCGCTGCACCCTAGGTC-CGACGCTGC-3' | fork restoration (FRET; leading strand gap) |

| | | |
|---|---|---|
| **#19** (62 nt) | 5'-CGTCGCAGCCTGGATCCCACGTCGCGAACAACTTCAGCTGATAGACACGTGGC-AATTGCCTA-(**Cy5**)-3' | |
| **#20** (30 nt) | 5'-GCAGCGTCGGACCTAGGGTGCAGCGCTTGT-3' | |
| **#21** (62 nt) | 5'-TAGGCATTTGCCGCGTGTGTATCACCTGAACATGTTCGCGACGTGGGATCCAGGCTG-CGACG-3' | |
| **#22** (62 nt) | 5'-GCAGCGTCGGACCTAGGGTGCAGCGCTTGT-3' | fork reversal from the four-way junction (FRET; leading strand gap) |
| **#23** (62 nt) | 5'-GGCAATTGCCACGTGTCTATCAGCTGAAGGACAAGCGCTGCACCCTAGGTCCGACG-CTGC-3' | |
| **#24** (62 nt) | 5'-(**Cy3**)-GCAGCGTCGGACCTAGGGTGCAGCGCTTGTTTTTCAGCTGATAGACACGTGGC-AATTGCC-3' | |
| **#25** (62 nt) | 5'-CCGTTAACGGTGCACAGATAGTCGACTTTTACAAGCGCTGCACCCTAGGTCCGA-CGCTGC-(**Cy5**)-3' | |

P1 – P1 nuclease sensitivity experiments; reversal – bulk SMARCAL1-mediated fork reversal experiment; MP – mass photometry; hom – homologous fork; het – heterologous fork

* Oligonucleotides used to assemble forks for cryo-EM analyses did not contain fluorescent labels.

**Supplemental Figure S1**:

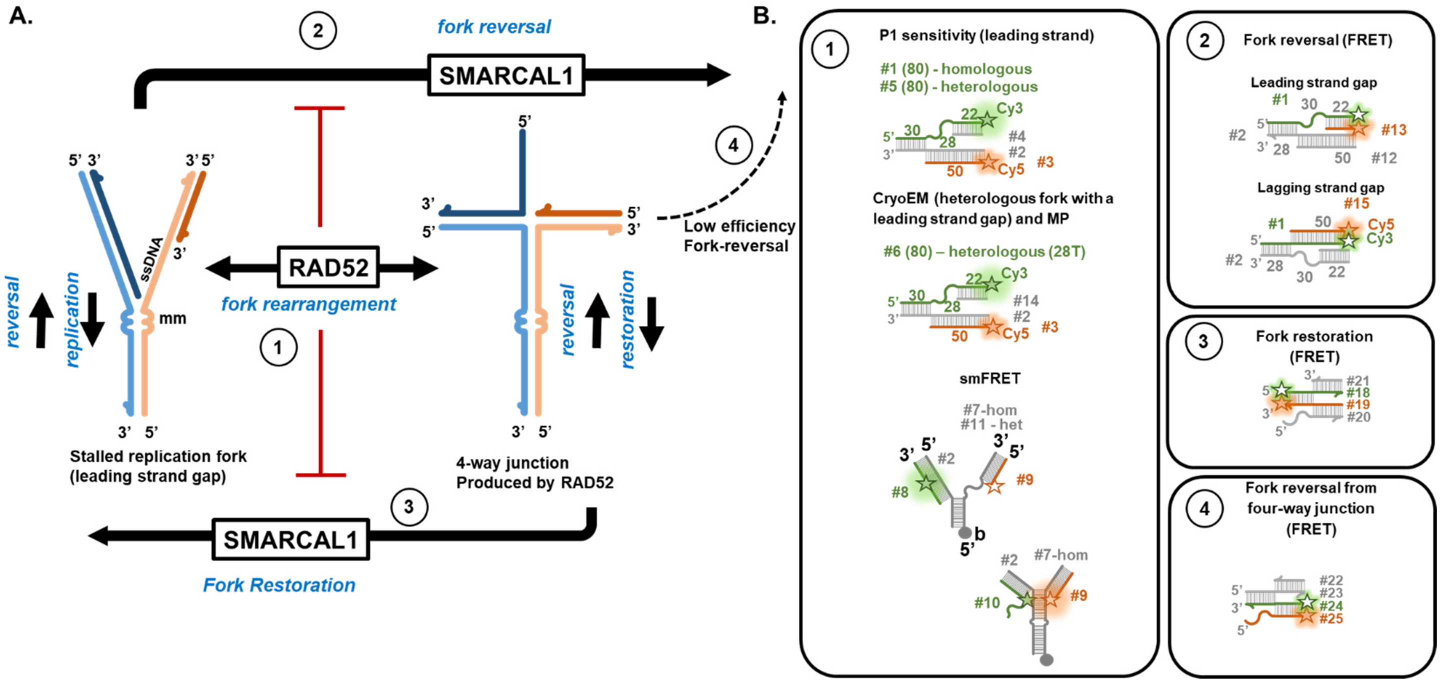


**Synthetic DNA structures mimicking stalled replication forks and intermediates of their processing used in this study. A**. Schematic of the replication fork rearrangement, reversal and restoration by RAD52 and SMARCAL1. **B.** Synthetic DNA structures representing each step schematically depicted in A. Each oligonucleotide number corresponds to a sequence listed in Table S1. Green and orange stars indicate Cy3 and Cy5 dyes, respectively.

**Supplemental Figure S2.**

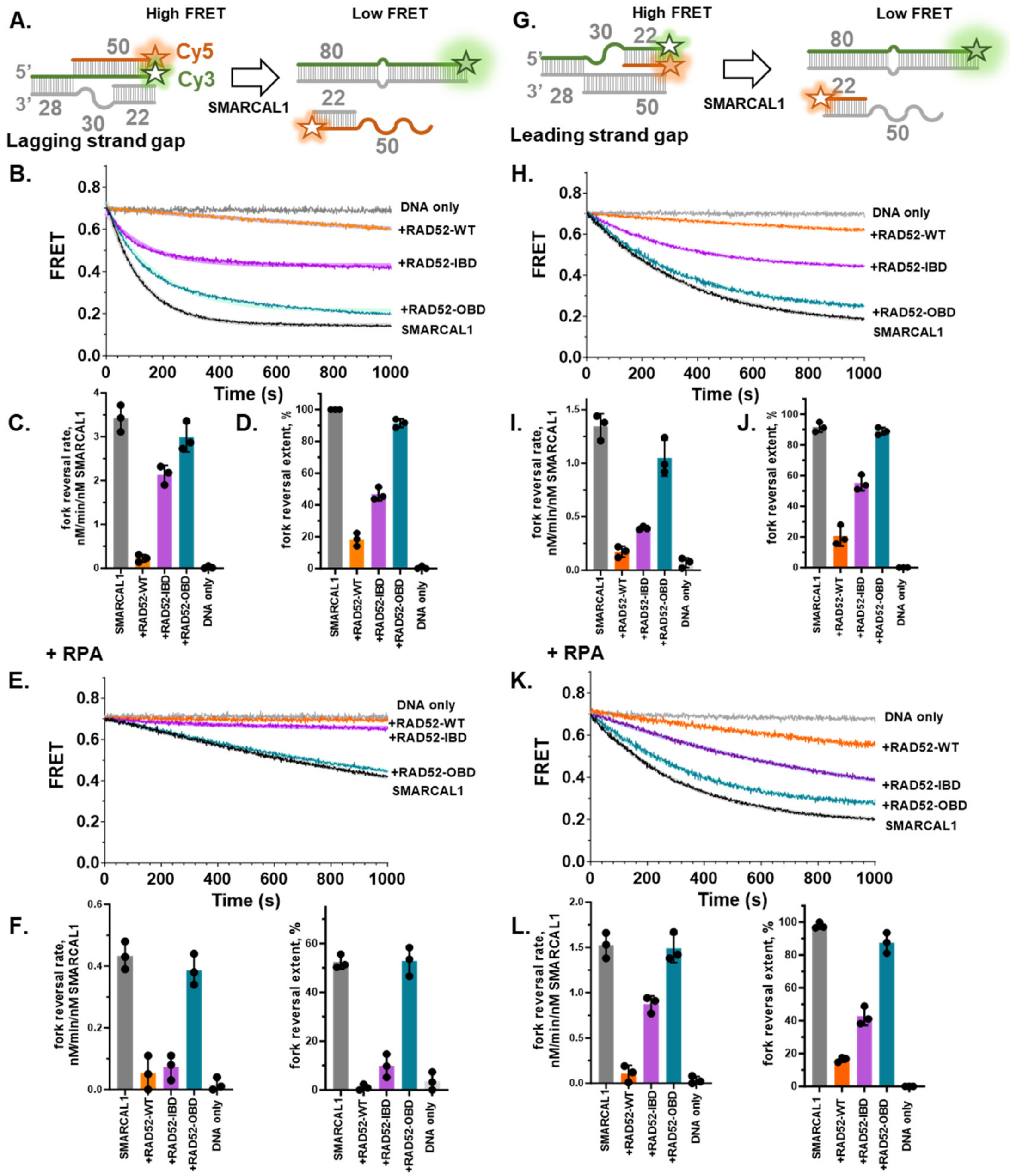


**The outer DNA binding site of RAD52 is more critical for fork protection that the inner binding site. A**. Experimental design of the FRET-based fork reversal experiment. Synthetic DNA structure mimicking stalled fork with a lagging strand gap was assembled from oligonucleotides #1, #2, #14 and #15 listed in **Supplemental Table S1** and schematically depicted in **Supplemental Figure S1** (manifold 2). The Cy3 (FRET donor) and Cy5 (FRET acceptor) are incorporated into the parental and nascent strands of the leading strand arm, respectively. Their proximity yields high FRET signal (~0.7). SMARCAL1-mediated fork reversal separates the two labeled oligonucleotides resulting in decrease in FRET to ~0.2. **B.** Representative time courses of the fork (5 nM) reversal by 0.5 nM SMARCAL1 in the absence (black) or presence of the wild type RAD52 (orange; 165 nM), $RAD52^{IBD}$ mutant (purple; 165 nM) or $RAD52^{OBD}$ mutant (teal; 165 nM). **C&D.** Quantification of rate (**C**) and extent (**D**) of the fork reversal reaction. Bar graphs represent the average and standard deviation for three independent experiments. **E-G** are the same as **B-D**, but in the presence of 10 nM RPA. **G-M** are the same as **A-F**, except for the fork containing the leading strand gap assembled from oligonucleotides #1, #2, #12 and #13. All data are plotted and analyzed in GraphPad Prism.

**Supplemental Figure S3:**

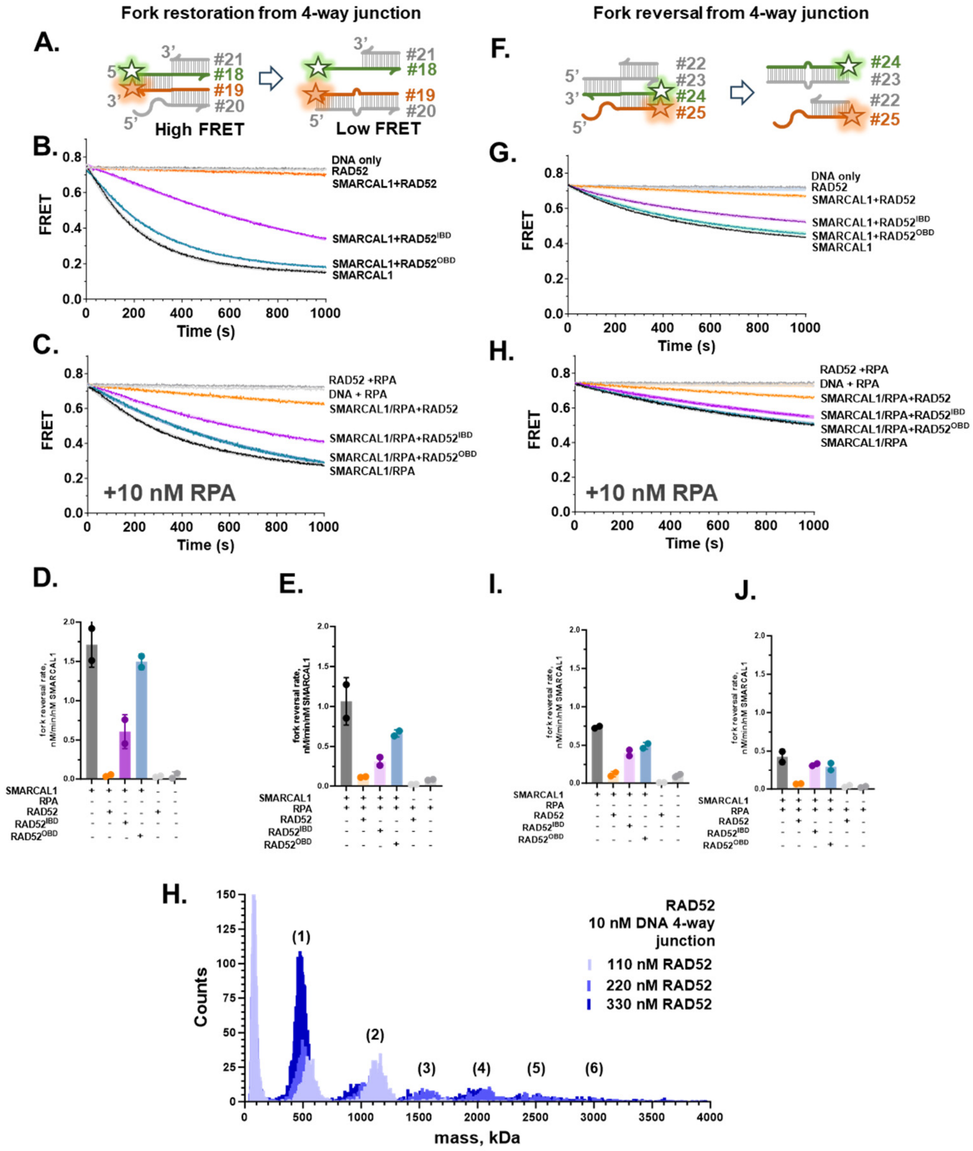


**SMARCAL1 efficiently restores but inefficiently reverses the fork remodeled by RAD52. The outer DNA binding site of RAD52 is critical for inhibition of SMARCAL1 activities on both of these substrates. A**. Experimental design of the FRET-based fork restoration experiment. Synthetic DNA structure mimicking four-way junction produced by the RAD52 activity on stalled fork with a leading strand gap was assembled from oligonucleotides #18-21 listed in **Supplemental Table S1** and schematically depicted in **Supplemental Figure S1** (manifold 3). The design of the substrates for fork restoration experiments was based on the Betous et al. paper[1]. Note that this substrate only allows fork remodeling in the restoration direction. The Cy3 (FRET donor) and Cy5 (FRET acceptor) are incorporated into the parental strands. Their proximity yields a high FRET signal (~0.7). SMARCAL1-mediated fork restoration separates the two labeled oligos resulting in decrease in FRET to ~0.2. **B.** Representative time courses of the fork (5 nM) reversal by 0.5 nM SMARCAL1 in the absence (black) or presence of the wild type RAD52 (orange; 165 nM), RAD52$^{IBD}$ mutant (purple; 165 nM) or RAD52$^{OBD}$ mutant (teal; 165 nM). **C** is the same as **B**, but in the presence of 10 nM RPA. **D&E.** Quantification of rate (**D**) and extent (**E**) of the fork restoration reaction. Bar graphs represent the average and standard deviation for three independent experiments. **F-J** are the same as **A-E**, but for the substrate that can proceed towards either restoration or reversal. The placement of labels in this substrate allows to specifically monitor fork reversal. **H.** Mass photometry experiment showing oligomeric states of 110, 220 and 330 nM RAD52 (from light to dark blue) bound four-way DNA junction substrate. Numbers above each peak indicate the number of RAD52 undecamers in each nucleoprotein complex.

**Supplemental Figure S4:**

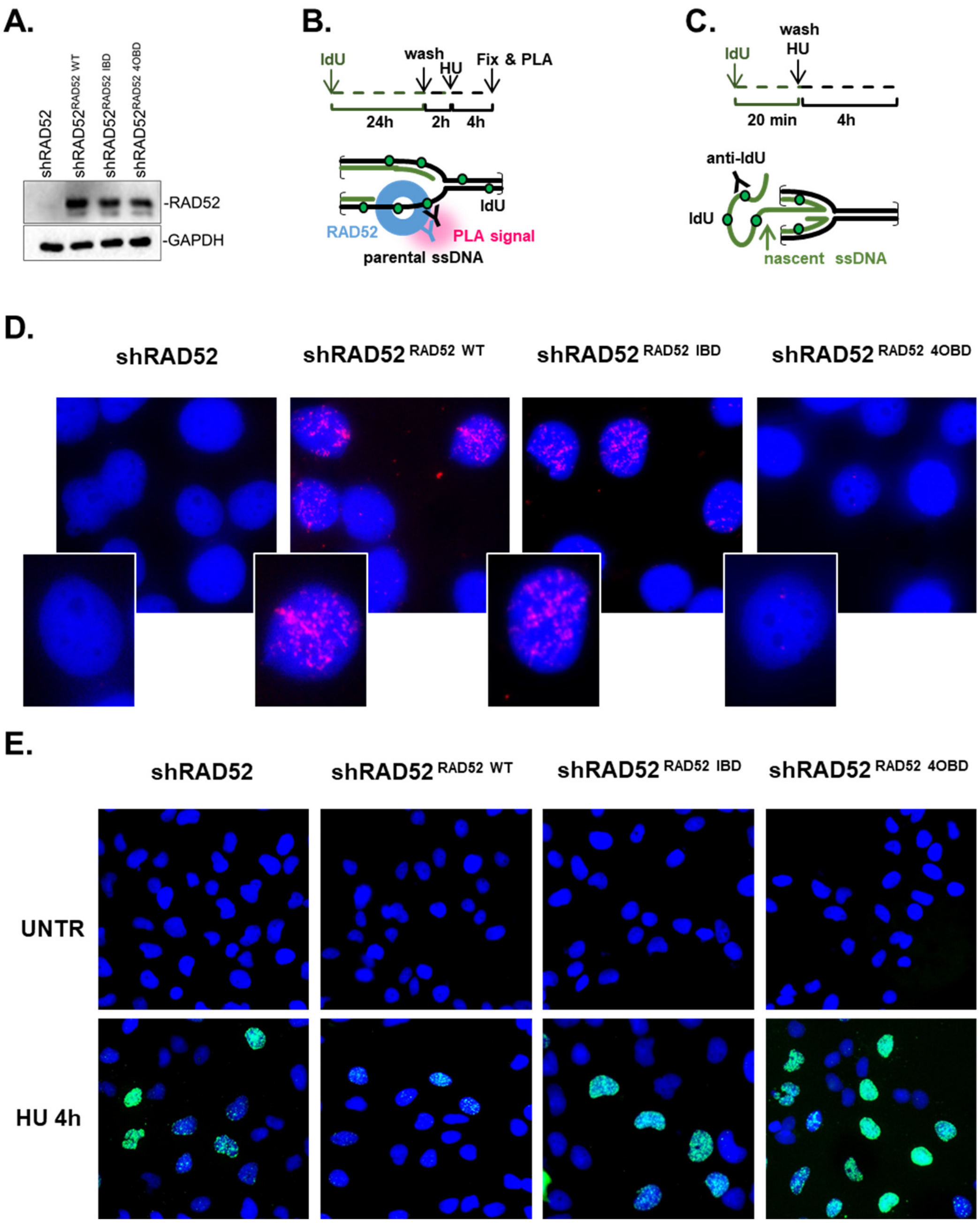


**Representative images used to quantify the cellular effects of RAD52 mutants shown in Fig 1I&J. A.** WB showing the expression level of RAD52, $RAD52^{IBD}$ and $RAD52^{OBD}$ in the transfected MRC5 cells depleted of endogenous RAD52. **B.** Schematic depiction of the PLA analysis of RAD52 binding to parental ssDNA. RAD52 knockdown MRC5 cells, complemented with the indicated RAD52 mutants were treated with 100 μM IdU for 20 hours, released for 2 hours in fresh medium and exposed to HU. The PLA reaction was carried out using antibodies against the RAD52 protein and IdU. **C.** Schematic depiction of the immunofluorescence analysis of the exposure of the nascent ssDNA upon replication fork reversal and degradation. RAD52 knockdown MRC5 cells complemented with the indicated RAD52 version were treated for 4h with HU. The ssDNA was detected by native anti-IdU immunofluorescence. **D.** Representative images from the PLA experiment showing that mutations in the inner (IBD) or outer (OBD) DNA binding sites of RAD52 differently affect RAD52 recruitment to parental gaps. Magnification of one nucleus is presented in each inset. **E.** Representative immunofluorescence images showing exposure of nascent ssDNA upon fork reversal and degradation. UNTR – cells untreated with HU.

**Supplemental Table S2: Summary for the Mass Photometry (MP) analysis of RAD52 and RAD52-fork DNA complexes**

| Molecular Species | | NH-Fork DNA | Monomer RAD52 | 1 ring (11mer) RAD52 | 1 ring RAD52-DNA | 2 rings RAD52-DNA | 3 rings RAD52-DNA |
|---|---|---|---|---|---|---|---|
| Theoretical mass (kDa) | | 75 | 48 | 528 | 603 | 1231 | 1659 |
| Experimental number and condition | Number of peaks | Mass (kDa) and peak population percentage | | | | | |
| 1. 10 nM NH-Fork DNA | 1 | 71±16.4 (100%) | | | | | |
| 2. 110 nM RAD52 | 2 | | 50±25 (11.8%) | 499±38 (88.2%) | | | |
| 3. 220 nM RAD52 | 2 | | 56±20 (5.7%) | 496±36 (94.3%) | | | |
| 4. 330 nM RAD52 | 2 | | 58±22 (5.2%) | 498±38 (94.8%) | | | |
| 5. 10 nM NH-Fork DNA+110 nM RAD52 | 3 | 62±18.4 (24.9%) | | 586±57 (70.4%) | | 1159±86 (4.7%) | |
| 6. 10 nM NH-Fork DNA+220 nM RAD52 | 4 | 53±17.6 (23.4%) | | 552±40 (49.6 %) | | 1077±63 (22.2%) | 1618±80 (4.8%) |
| 7. 10 nM NH-Fork DNA+330 nM RAD52 | 4 | 60±19.6 (32.7%) | | 546±43 (13.0%) | | 1060±53 (48.0 %) | 1060±53 (6.3 %) |
| 8. 10 nM NH-Fork DNA | 1 | 71±13.0 (100%) | | | | | |
| 9. 110 nM RAD52-IBD | 1 | | | 441±34 (100%) | | | |
| 10. 220 nM RAD52-IBD | 1 | | | 440±34 (100%) | | | |
| 11. 330 nM RAD52-IBD | 1 | | | 444±38 (100%) | | | |
| 12. 10 nM NH-Fork DNA+110 nM RAD52-IBD | 3 | 79±21.0 (45.4%) | | 487±54 (50.5%) | | 953±64 (3.9%) | |
| 13. 10 nM NH-Fork DNA+220 nM RAD52-IBD | 3 | 79±17.6 (12.7%) | | 495±50 (79.0 %) | | 964±77 (8.3%) | |
| 14. 10 nM NH-Fork DNA+330 nM RAD52-IBD | 3 | 85±17.7 (5.9%) | | 492±50 (80.0%) | | 940±71 (14.1 %) | |
| 15. 10 nM NH-Fork DNA | 1 | 76±22.0 (100%) | | | | | |
| 16. 110 nM RAD52-OBD | 2 | | 66±34 (11.7%) | 464±47 (88.3%) | | | |
| 17. 220 nM RAD52-OBD | 2 | | 72±28 (9.3%) | 456±53 (90.7%) | | | |
| 18. 330 nM RAD52-OBD | 2 | | 96±41 (5.5%) | 464±52 (94.5%) | | | |
| 19. 10 nM NH-Fork DNA+110 nM RAD52-IBD | 2 | 92±23 (54.6%) | | 469±58 (45.4%) | | | |
| 20. 10 nM NH-Fork DNA+220 nM RAD52-IBD | 2 | 93±27 (33.3%) | | 477±56 (66.7 %) | | | |
| 21. 10 nM NH-Fork DNA+330 nM RAD52-IBD | 2 | 79±23 (15.5%) | | 497±53 (84.5%) | | | |

The data in this table are representative of three independent experiments. All data were analyzed using DiscoverMP software package. The molecular weights correspond to the mean (± standard deviation) of respective Gaussians peaks. The percentage of the counts in each peak is shown in the parenthesis.

**Supplemental Figure S5:**

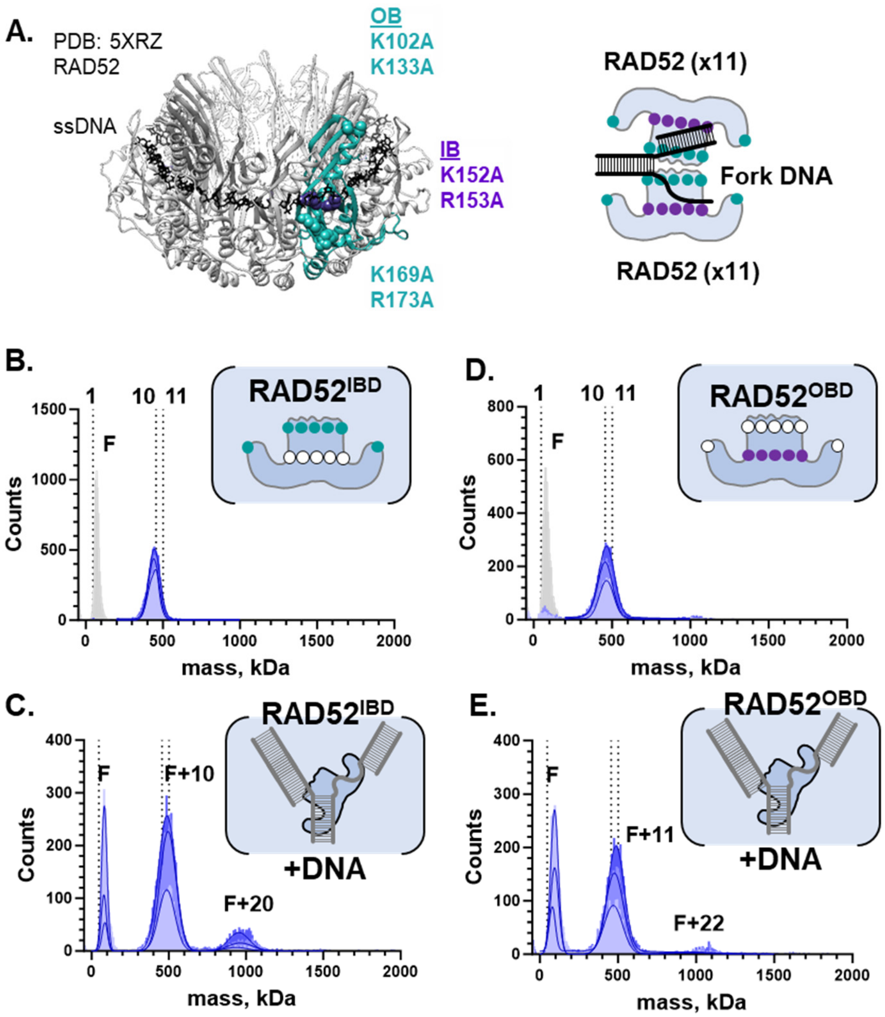


**Mutations in the DNA binding sites of RAD52 affect the formation of the double-ring RAD52-fork complex.** **A.** Location of the inner DNA binding site (K152/R153; purple) and the bipartite outer DNA binding site (K102/K133/K169/R173) in one of the monomers (teal ribbon representation) in an undecamerc RAD52 ring. The mutations are mapped on the crystal structure of the RAD52-ssDNA complex (PDB: 5XRZ). Cartoon on the right shows position of the two binding site within the double-ring RAD52 structure. **B-E.** Mass photometry (MP) analysis of the macromolecular complexes formed by the mutant forms of RAD52. In all experiments, representative distributions are shown for 100 nM (11xFork; light blue), 220 nM (22xFork; medium blue), and 330 nM (33xFork; dark blue) $RAD52^{IBD}$ or $RAD52^{OBD}$. Lines correspond to fitting of the molecular weights distributions with multiple Gaussians using GrapPad Prism. Vertical dotted lines indicate molecular weights of RAd52 monomer, decamer and undecamer, respectively. Quantification of the peaks performed in the DiscoverMP software is detailed in the **Supplemental Table S2**. A heterologous DNA fork with a 30 nt lagging strand gap was assembled using oligonucleotides #2, #3, #4 and #6 listed in the **Supplemental Table S1. B.** Molecular weight distribution of the $RAD52^{IBD}$ in solution. Note that the $RAD52^{IBD}$ forms lower molecular weight complexes than the wild type RAD52. **C.** Molecular weight distributions of the $RAD52^{IBD}$ bound to heterologous fork. While binding to the fork is evident from the decrease in the peak corresponding for free fork and a shift of the peak corresponding to the $RAD52^{IBD}$ single ring, the amounts of double-ring complexes are lower compared to the wild type RAD52. **D.** Molecular weight distribution of the $RAD52^{OBD}$ in solution. **E.** Molecular weight distributions of the $RAD52^{OBD}$ bound to heterologous fork. Virtually no double rings are observed with this mutant. The shift in the single ring peak relative to the free $RAD52^{OBD}$ suggests binding to the fork.

**Supplemental Figure S6:**

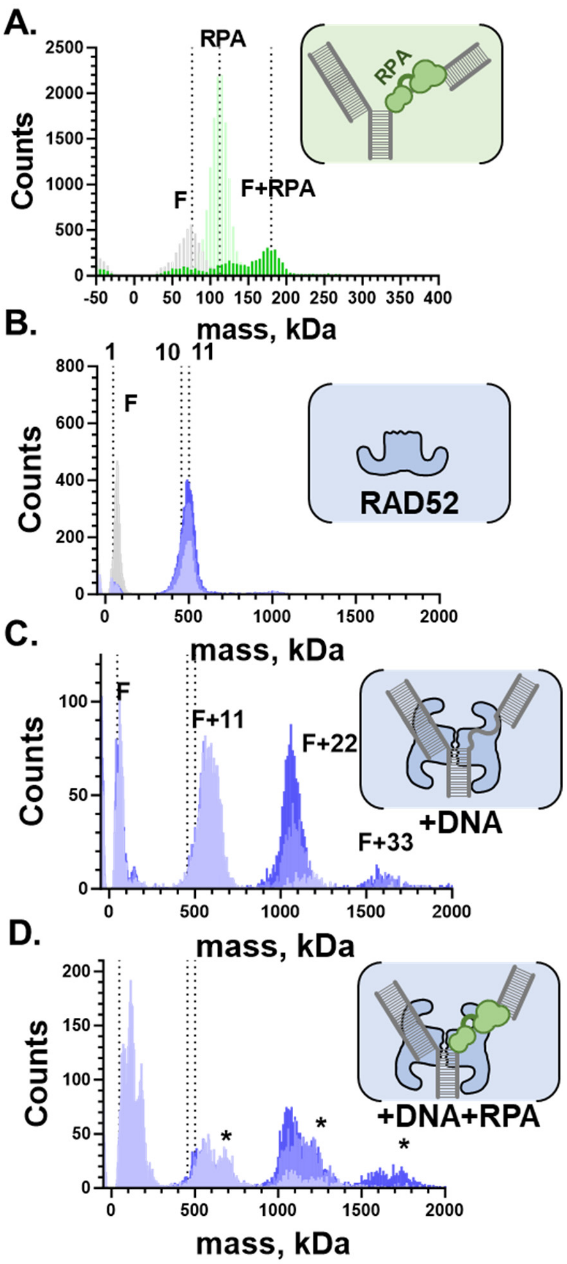


**RAD52 forms double-undecamer structures on the model fork bound by RPA. A.** The MP analysis of the 10 nM DNA fork (grey; the heterologous fork with a 30 nt lagging strand gap was assembled using oligonucleotides #2, #3, #4 and #6 listed in the **Supplemental Table S1**), 10 nM RPA (light green) and the fork-RPA complex (dark green). **B.** The MP analysis of RAD52. **C.** RAD52 + plus 10 nM fork. **D.** Fork DNA bound by RPA and RAD52. In all experiments, representative distributions are shown for 100 nM (11xFork; light blue), 220 nM (22xFork; medium blue), and 330 nM (33xFork; dark blue) RAD52. The RAD52-fork-RPA complexes are marked with * in **D**.

**Supplemental Table S3: Summary for the Mass Photometry (MP) analysis of the RPA and RAD52-RPA-fork DNA complexes**

| Moleculer Species | | Monomer RAD52 | (het)Fork DNA | RPA | RPA-DNA | 1 ring RAD52 (-DNA) | 1 ring RAD52 (-DNA)-RPA | 2 rings RAD52-DNA | 2 rings RAD52-DNA-RPA | 3 rings RAD52-DNA | 3 rings RAD52-DNA-RPA |
|---|---|---|---|---|---|---|---|---|---|---|---|
| Theoretical mass (kDa) | | 48 | 75 | 111 | 186 | 528-603 | 639-714 | 1231 | 1342 | 1659 | 1806 |
| Experimental number and condition | Number of peaks | | Mass (kDa) and peak population percentage | | | | | | | | |
| 1. 10 nM (het)Fork DNA | 1 | | 68±13.6 (100%) | | | | | | | | |
| 2. 10 nM RPA | 1 | | | 112±11 (100%) | | | | | | | |
| 3. 10 nM (het)Fork DNA+10 nM RPA | 3 | | 65±22.1 (19.8%) | 124±24 (34.5%) | 176±16 (45.7%) | | | | | | |
| 4. 10 nM (het)Fork DNA+10 nM RPA+110 nM RAD52 | 7 | 69±17.5 (18.6%) | | 117±19 (28.5%) | 173±23 (19.3%) | 562±46 (15.8%) | 674±47 (11.9%) | 1075±52 (3.3%) | 1168±85 (2.6%) | | |
| 5. 10 nM (het)Fork DNA+10 nMRPA+220 nM RAD52 | 6 | 77±35.0 (16.0%) | | 114±25 (26.7%) | | 554±53 (15.5%) | 668±54 (10.6%) | 1063±54 (10.5%) | 1154±102 (20.5%) | | |
| 6. 10 nM (het)Fork DNA+10 nM RPA+330 nM RAD52 | 6 | 75±17.4 (13.1%) | | 111±31 (26.5%) | | 522±47 (13.2 %) | | 1069±61 (21.1%) | 1183±67 (16.3%) | 1596±74 (5.6%) | 1720±85 (4.1%) |
| 7. 10 nM RPA+110 nM RAD52 | 3 | 46±9.2 (11.6%) | | 122±14 (68.7%) | | 546±34 (19.8%) | | | | | |
| 8. 10 nMRPA+220 nM RAD52 | 3 | 57±13.6 (6.1%) | | 123±19 (34.8%) | | 552±39 (59.0%) | | | | | |
| 9. 10 nM RPA+330 nM RAD52 | 2 | | | 125±17 (16.3%) | | 549±43 (83.7 %) | | | | | |

The data in this table are representative of three independent experiments. All data were analyzed using DiscoverMP software package. The molecular weights correspond to the average (± standard deviation) of respective Gaussians peaks. The percentage of the counts in each peak is shown in the parenthesis. (het)Fork – heterologous fork annealed using oligonucleotides #1, #2, #3 and #.

### Supplemental Table 4: Cryo-EM data collection

| **Structure** | **apoRAD52** | **RAD52-fork DNA (J369)** | **RAD52-fork DNA (J209)** | **RAD52-fork DNA (J212)** |
|---|---|---|---|---|
| EMDB accession | EMD-41357 | EMD-42440 | EMD-42066 | EMD-42069 |
| PDB code | 8TKQ | - | - | - |
| **Data collection and image processing** | | | | |
| Microscope | Titan Krios | Titan Krios | Titan Krios | Titan Krios |
| Voltage (kV) | 300 | 300 | 300 | 300 |
| Camera | TFS Krios | Falcon IV | Falcon IV | Falcon IV |
| Magnification | 130,000 x | 130,000 x | 130,000 x | 130,000 x |
| Electron exposure ($e^-$/Å$^2$) | 50 | 56 | 56 | 56 |
| Exposure time (s) | 1.8 | 7 | 7 | 7 |
| Number of frames | 50 | - | - | - |
| Defocus range (mm) | -1.0 to -2.5 | -0.8 to -2.5 | -0.8 to -2.5 | -0.8 to -2.5 |
| Pixel size (Å) | 0.8255 | 0.959 | 0.959 | 0.959 |
| Number of movies | 11,290 | 6,093 | 6,093 | 6,093 |
| Particles for final reconstruction (no.) | 623,559 | 124,067 | 80,665 | 33,883 |
| Map resolution (Å) | 2.5 | 6.75 | 5.35 | 8.5 |
| Half map FSC threshold | 0.143 | 0.143 | 0.143 | 0.143 |

**Supplemental Figure S7:**

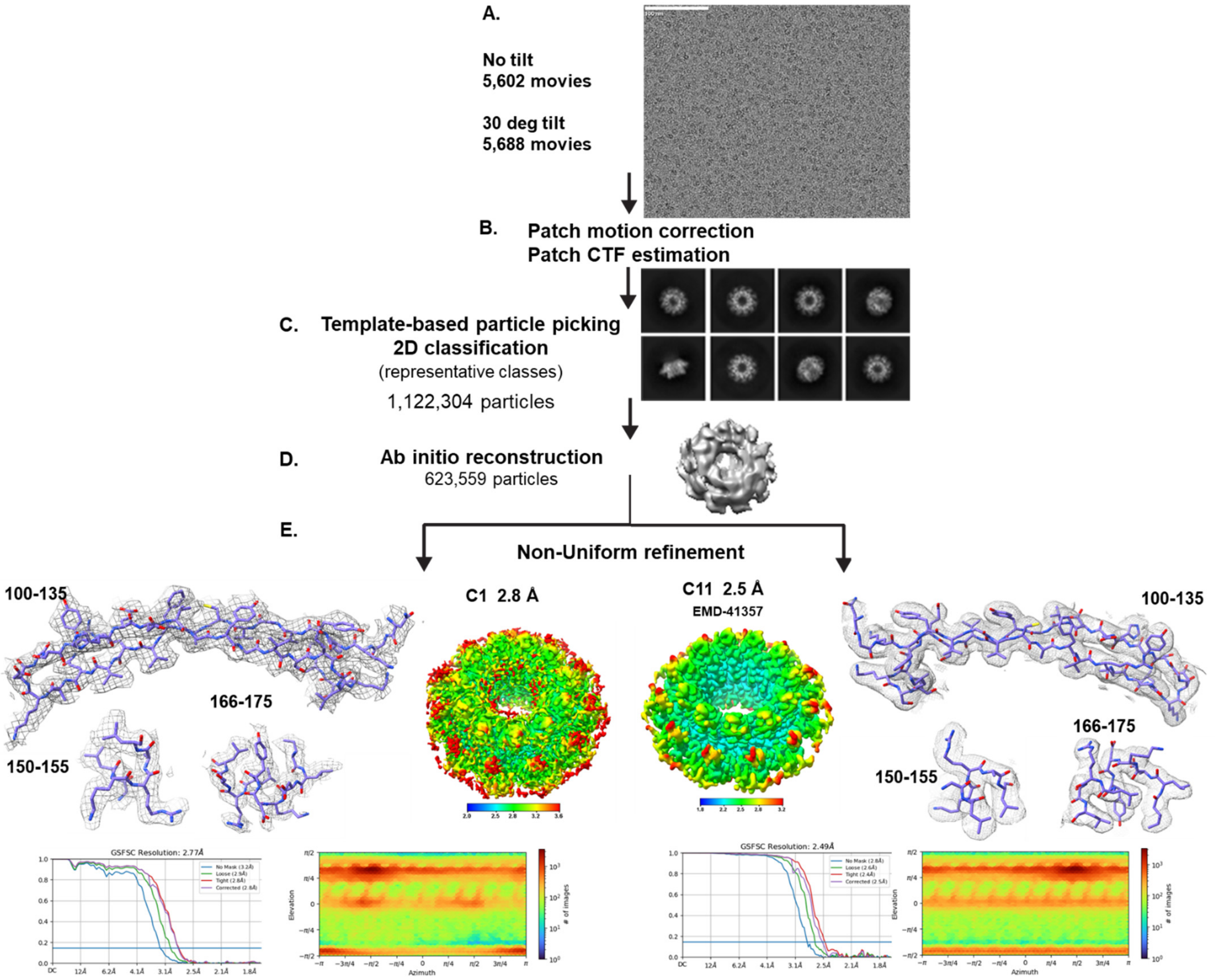


**Workflow for obtaining the cryo-EM structure of the RAD52 undecamer. A.** Two data sets, 30° tilted and non-tilted, were collected for apo RAD52. **B-C.** For single particle analysis, micrographs were manually curated and multiple rounds of 2D classification were performed yielding a final stack of 623,559 particles which were used for *ab-initio* 3D reconstruction (**D.**). **E.** The final structures of apo RAD52 were obtained by performing a non-uniform refinement by imposing C1 (2.8Å, left) and C11 (2.5Å, right; EMD-41357) symmetries. The residues in the structures are colored by resolution suggesting the rigid core and DNA binding regions (blue) and more flexible loops (red). The three regions containing the DNA binding residues are shown as zoom-ins. The heatmaps of the angular distribution of particles used to generate the final structures of apo RAD52 are shown.

**Supplemental Figure S8:**

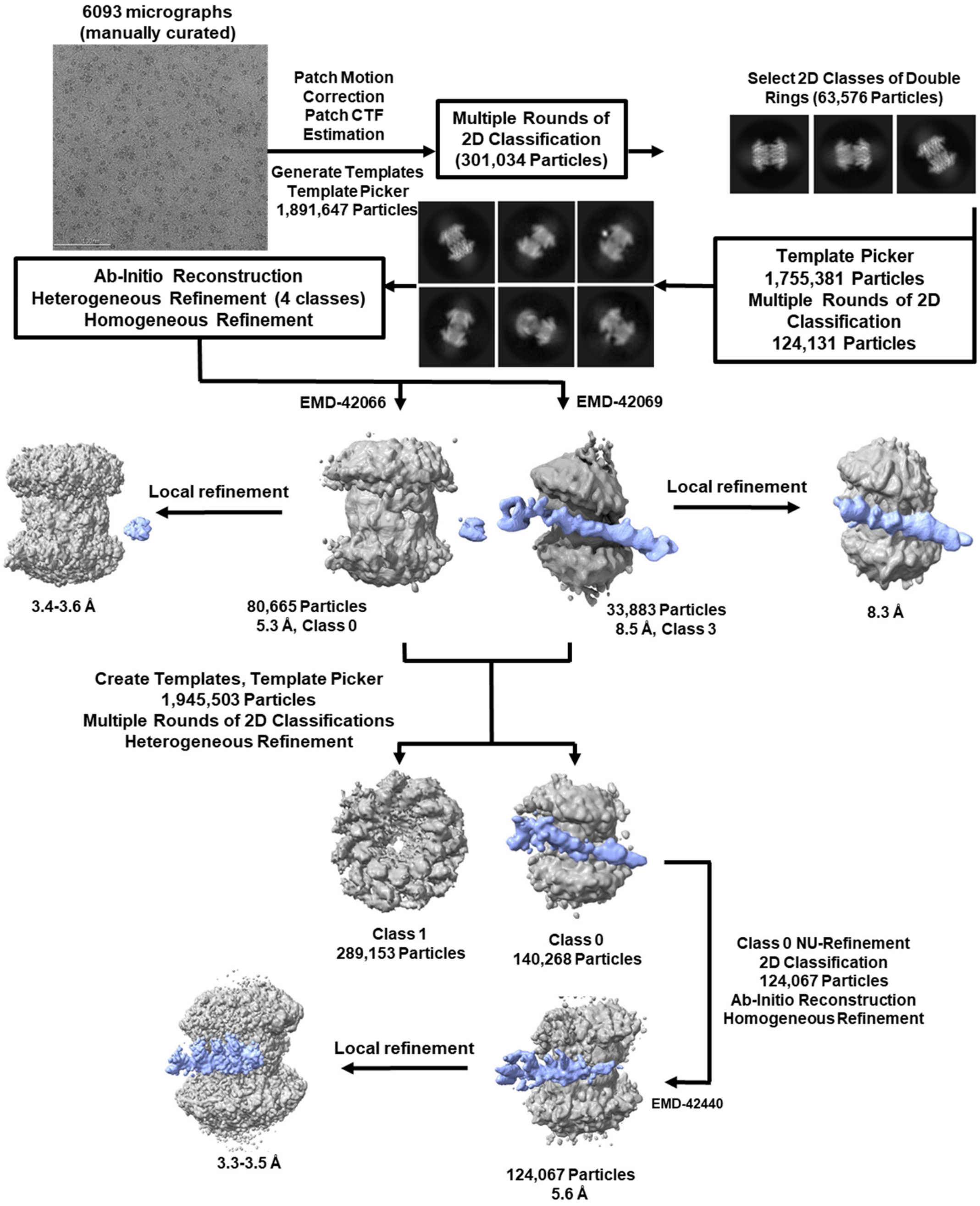


**Workflow for obtaining the cryo-EM structures of the RAD52 double undecamer bound to synthetic fork DNA.** For single particle analysis, micrographs were manually curated and multiple rounds of 2D classification performed yielding a final stack of 63,576 particles of 2D classes containing RAD52 double rings. The selected 2D class averages were used as templates followed by multiple rounds of 2D classification yielding 124,131 particles prior to *ab-initio* reconstruction, heterogeneous refinement (4 classes) and homogeneous refinement of final structures of RAD52-fork DNA. The heatmaps of the angular distribution of particles used to generate the final RAD52-fork DNA structures (EMD-42066 and EMD-42069) are shown. The templates were then created from final maps and used for template picking and particle extraction from micrographs (1,945,503 particles) followed by multiple rounds of 2D classification and heterogeneous refinement. The final double-ring structure of EMD-42440 was obtained by performing non-uniform refinement, 2D classification, *ab-initio* reconstruction, and homogeneous refinement. The local refinement of each ring (top ring and bottom ring with fork DNA) was carried out without particle subtraction. The masks were generated for the top ring and then for the bottom ring with fork DNA, which were used for local refinement in cryoSPARC. The maps from local refinement jobs were combined using ChimeraX.

**Supplemental Figure S9:**

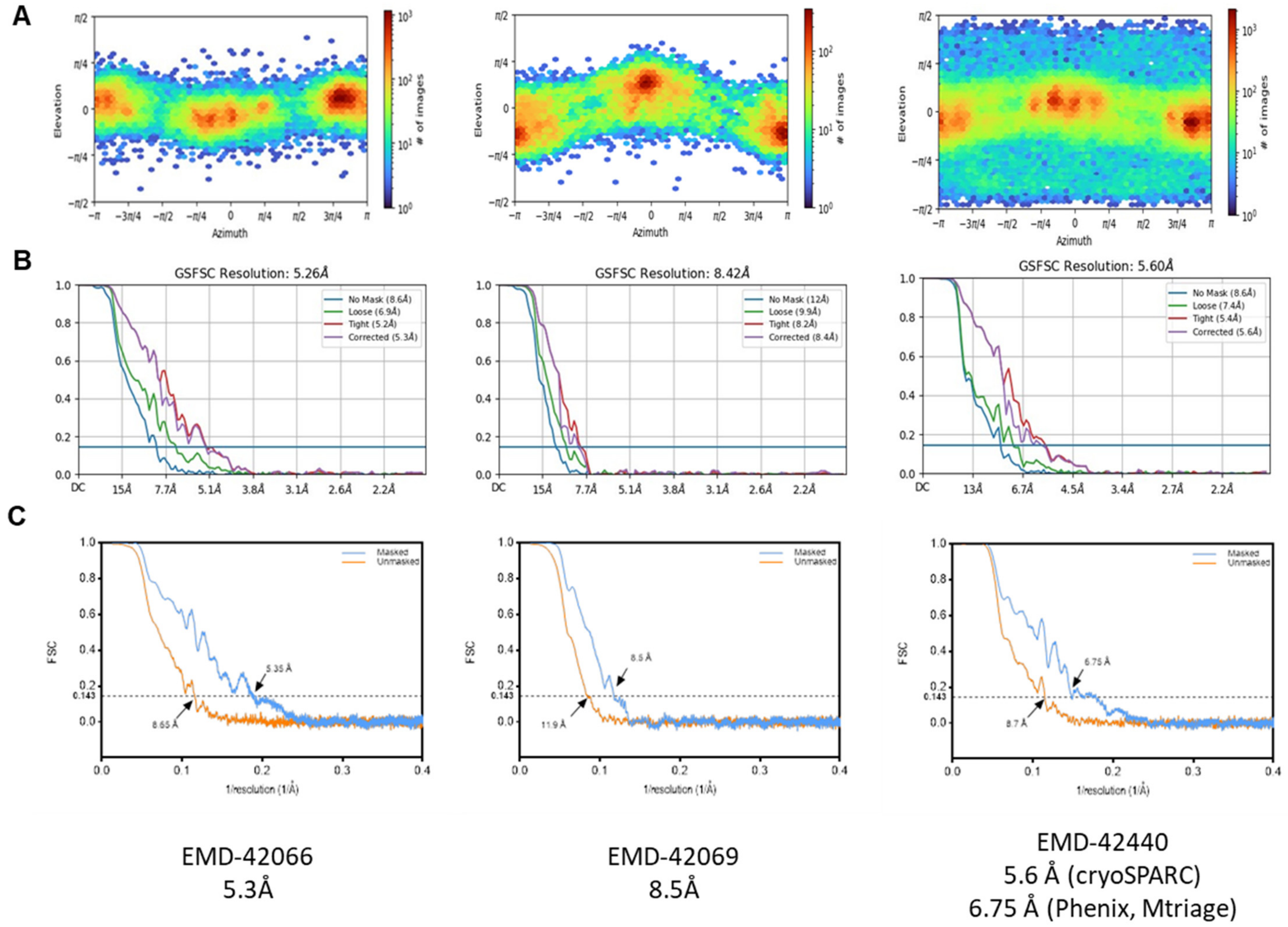


**The heatmaps and resolution evaluation for the RAD52-fork structures. (A)** The heatmap of the angular distribution of particles used to generate the final RAD52-fork DNA structures are shown. **(B)** GSFSC curves from cryoSPARC are shown for the final RAD52-fork DNA structures. **(C)** FSC curves with and without mask were calculated using Mtriage as part of Phenix package.

**Supplemental Figure S10.**

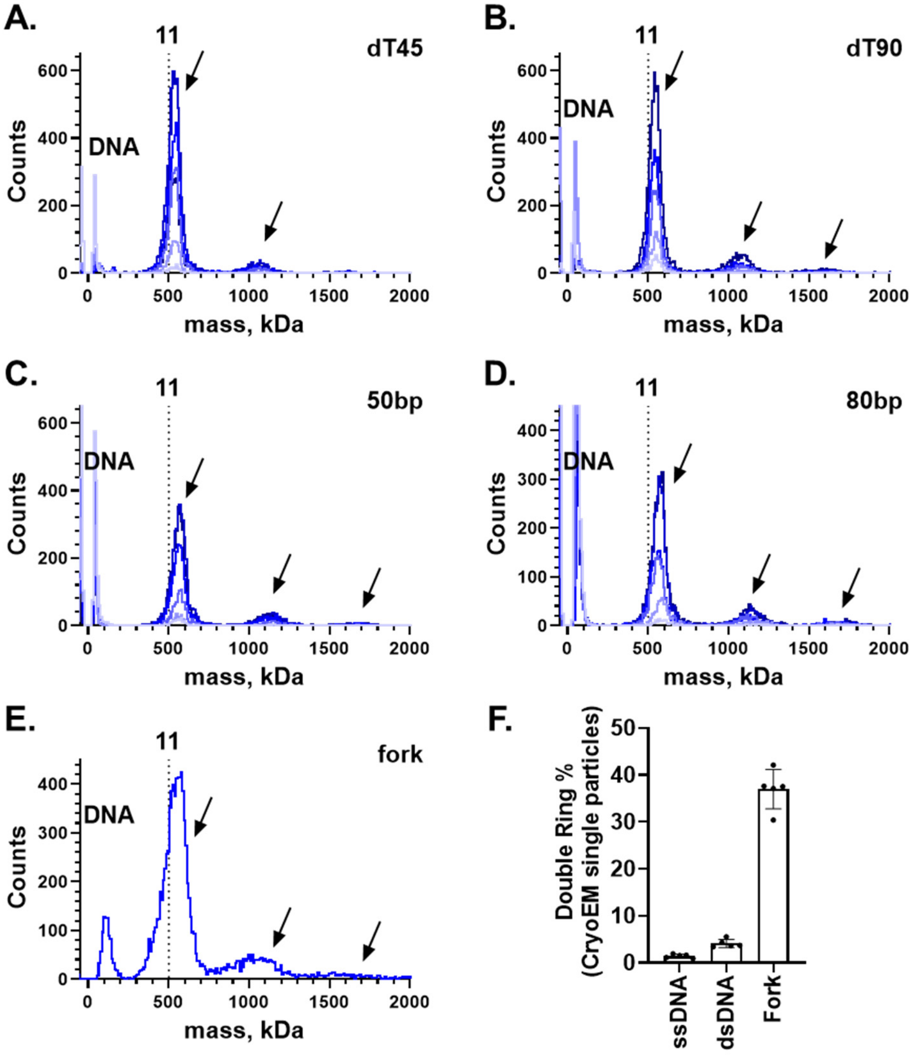


**The spool-like arrangement of the RAD52 double rings is specific to the replication fork structure. A-D.** Mass photometry experiments showing oligomeric states of RAD52 bound to dT45 ssDNA (**A**), dT90 ssDNA (**B**), 50 bp dsDNA (annealed using oligonucleotides #12 and #15; **C**) and 80 bp dsDNA (annealed using oligonucleotides #1 and #2; **D**). The ratio of RAD52 monomers to DNA was 7x, 11x, 22x, 33x, and 44x (from light to dark blue). In each experiment, RAD52 was incubated with 200 nM (molecules) of the respective DNA substrate in the presence of 2 mM BS3 cross-linker. The reaction mixture was then diluted 40-fold into the imaging buffer. Note that these are the same conditions that were used in sample preparation for CryoEM. Arrows indicate positions of single, double and triple RAD52 undecamer bound to DNA. The vertical line indicates the mass of unbound RAD52 undecamer. **E**. Five μM RAD52 (25x) was incubated with 200 nM non-homologous fork DNA in the presence of 2 mM BS3 cross-linker. The sample was then diluted 20-fold into the imaging buffer. **F**. Quantification of particles from the cryo-EM analysis of the fork DNA, ssDNA (dT90) and dsDNA (80 bp). All data were collected at using 5 μM RAD52 (25x) incubated with 200 nM of respective DNA in the presence of BS3. Each point corresponds to a different particle selection/sorting and represents the number of particles in all two-ring classes divided by total number of picked particles. While in the case of replication fork DNA ~40% of particles were found in the in the classes containing a spool-like arrangement of the double-rings, only few particles with such architectures were observed when RAD52 was bound to ssDNA or dsDNA substrates.

**Supplemental Figure S11:**

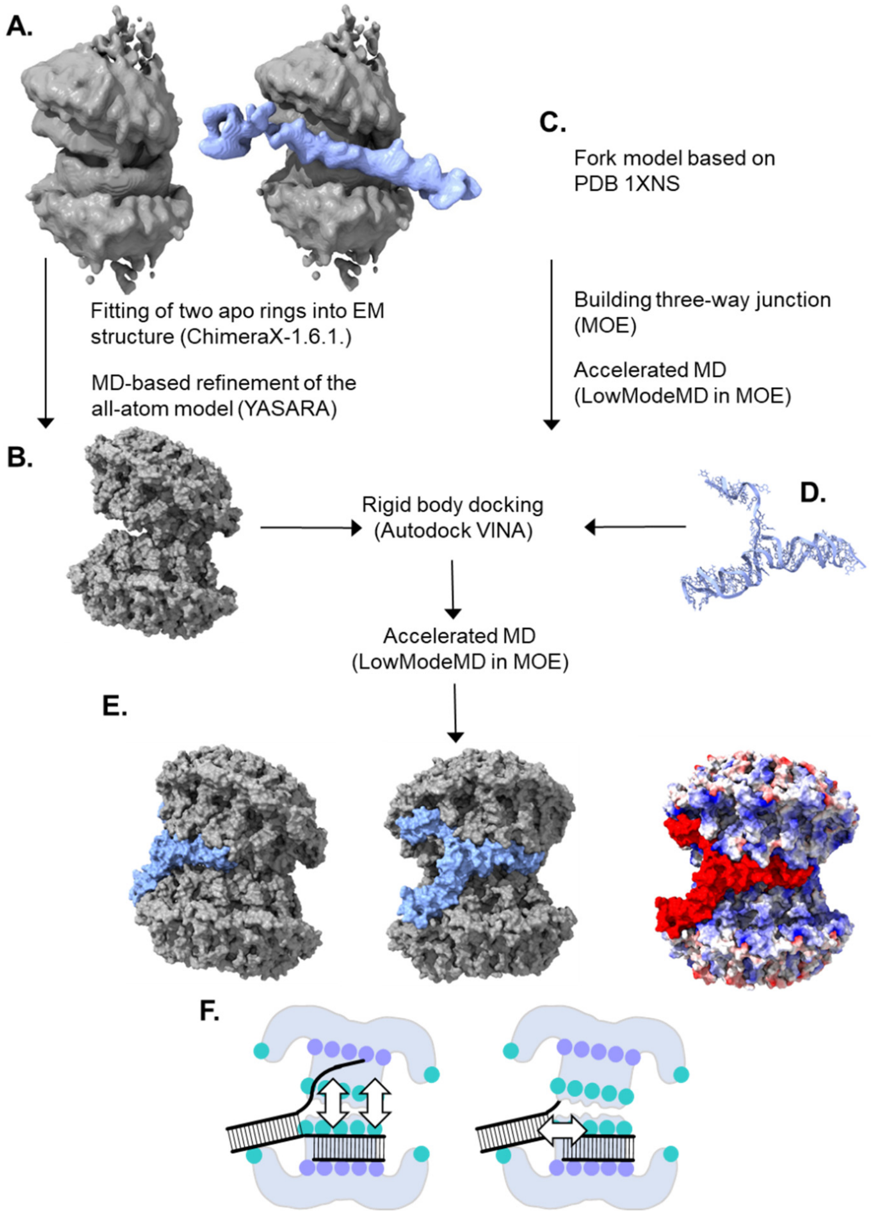


**Computational modeling workflow**. **A.&B**. The cryo-EM density map of EMD-42069 was used for placement of the two undecameric RAD52 rings from the apo structure (PDB 8TKQ). The resulting structure was refined using YASARA knowledge-based force field and a simulated annealing molecular dynamics (MD) protocol. **C.&D**. A small fork DNA structure was produced from the four-way junction in PDB 1XNS by retaining the oligonucleotides #16-17 (see **Supplemental Table S1**) using accelerated MD protocol (see Methods). **E.** Model developed by docking of the refined model fork structure into all-atom structure of the RAD52 double ring followed by another round of accelerated MD. Two different views show the extent of the complex between RAD52 (grey) and the fork (blue). On the right, the RAD52 structure is colored based on electrostatics. **F.** Two possible mechanisms for the RAD52-mediated strand exchange reaction at the fork. The dynamic exchange may proceed on the surface of the spool-like double-ring arrangement of RAD52 (left). Alternatively, the fork can split at the barrel (right).

**Supplemental Figure S12:**

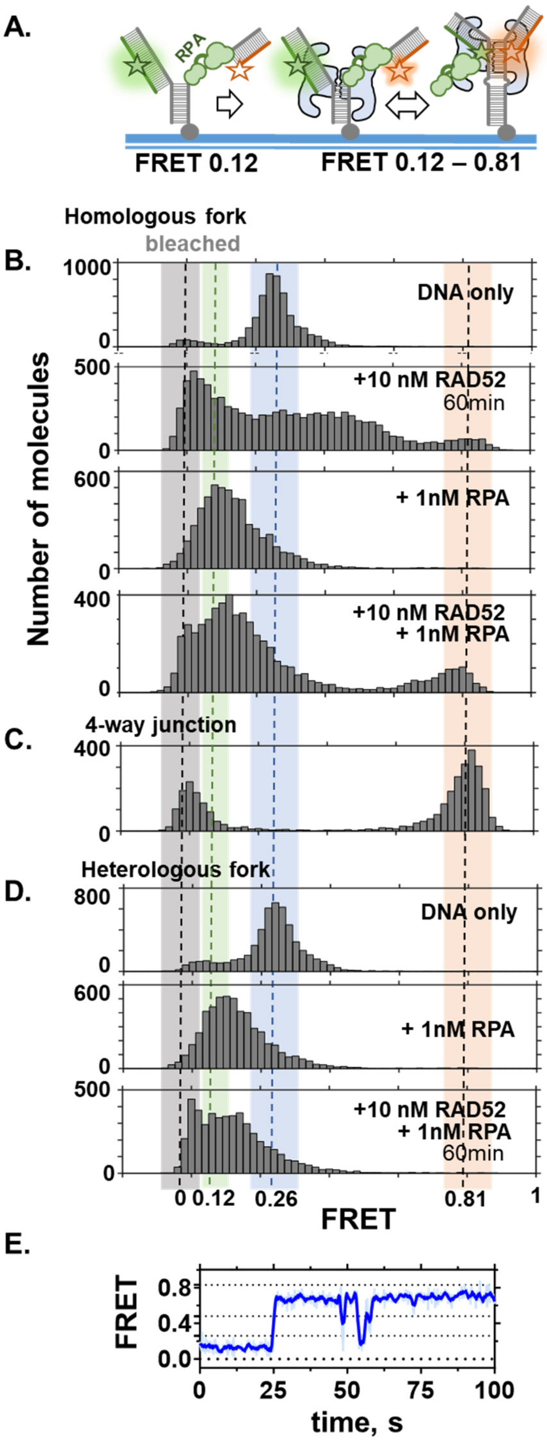


**RPA constrains the dynamics of the RAD52-mediated DNA strand exchange at the fork, but does not reduce its efficiency. A.** Cartoon depiction of the experimental design. A model replication fork with a leading strand gap was assembled from oligonucleotides #2, #7, #8 and #9 (homologous fork), #2, #7, #9 and #10 (four-way junction), or #2, #8, #9 and #11 (heterologous fork) listed in **Supplemental Table S1**. The Cy3 (FRET donor) and Cy5 (FRET acceptor) dyes are placed at the lagging and leading arms of the fork (FRET 0.26). **B-D.** Single-molecule FRET distributions for the forks containing two fully homologous arms (homologous fork) (**B**), fully exchanged fork (4-way junction) (**C**), and heterologous fork (the ssDNA gap region consists of 30 Ts) (**D**). Note that the peak around 0 FRET corresponds to the molecules with bleached Cy5 dye which accumulate over time. **E**. A representative smFRET trajectory for the RAD52-mediated strand exchange reaction in the presence of RPA.

**Supplemental Table 5**: **Summary for the Mass Photometry (MP) analysis of the SMARCAL1- and RAD52-fork DNA complexes**

<table>
<tr><td>Molecular Species</td><td></td><td>NH-Fork DNA</td><td>SMARCAL1</td><td>SMARCAL1-DNA</td><td>1 ring RAD52</td><td>1 ring RAD52-DNA</td><td>2 rings RAD52-DNA</td><td>3 rings RAD52-DNA</td></tr>
<tr><td>Theoretical mass (kDa)</td><td></td><td>75</td><td>107</td><td>182</td><td>528</td><td>603</td><td>1231</td><td>1659</td></tr>
<tr><td>Heterologous Fork</td><td>Number of peaks</td><td colspan="7">Mass (kDa) and peak population percentage</td></tr>
<tr><td>1. 10 nM NH-Fork DNA</td><td>1</td><td>70±14.6 (100%)</td><td></td><td></td><td></td><td></td><td></td><td></td></tr>
<tr><td>2. 10 nM SMARCAL1</td><td>1</td><td></td><td>99±23 (100%)</td><td></td><td></td><td></td><td></td><td></td></tr>
<tr><td>3. 10 nM NH-Fork DNA+10 nM SMARCAL1</td><td>2</td><td>70±16.1 (63.9%)</td><td></td><td>167±17 (36.1%)</td><td></td><td></td><td></td><td></td></tr>
<tr><td>4. 10 nM NH-Fork DNA+10 nM SMARCAL1+110 nM RAD52</td><td>4</td><td colspan="2">70±19.6 (29.3%)</td><td>168±20 (18.2%)</td><td colspan="2">556±48 (28.5%)<br>614±35 (20.6%)*</td><td>1115±87 (3.5%)</td><td></td></tr>
<tr><td>5. 10 nM NH-Fork DNA+10 nM SMARCAL1+220 nM RAD52</td><td>4</td><td colspan="2">72±31.5 (26.5%)</td><td></td><td colspan="2">551±53 (27.6%)</td><td>1160±82 (40.5%)</td><td>1597±90 (5.4%)</td></tr>
<tr><td>6. 10 nM NH-Fork DNA+10 nM SMARCAL1+330 nM RAD52</td><td>4</td><td colspan="2">77±35.0 (30.5%)</td><td></td><td colspan="2">489±56 (9.3%)</td><td>1063±53 (45.8%)</td><td>1577±69 (14.4%)</td></tr>
<tr><td>7. 10 nM NH-Fork DNA+10 nM SMARCAL1+110 nM RAD52-IBD</td><td>4</td><td colspan="2">73±14.3 (54.0%)</td><td>164±33 (9.8%)</td><td colspan="2">508±42 (29.5%)</td><td>976±60 (6.8%)</td><td></td></tr>
<tr><td>8. 10 nM NH-Fork DNA+10 nM SMARCAL1+220 nM RAD52-IBD</td><td>4</td><td colspan="2">83±24.0 (11.1%)</td><td></td><td colspan="2">450±43 (66.1%)</td><td>962±60 (13.1%)</td><td>1423±70 (9.7%)</td></tr>
<tr><td>9. 10 nM NH-Fork DNA+10 nM SMARCAL1+330 nM RAD52-IBD</td><td>4</td><td colspan="2">78±21.2 (17.7%)</td><td></td><td colspan="2">443±45 (60.4 %)</td><td>974±69 (11.9%)</td><td>14224±54 (9.9%)</td></tr>
<tr><td>10. 10 nM NH-Fork DNA+10 nM SMARCAL1+110 nM RAD52-OBD</td><td>3</td><td colspan="2">70±17.3 (6.8%)</td><td>167±20 (9.3%)</td><td colspan="2">487±53 (83.9%)</td><td></td><td></td></tr>
<tr><td>11. 10 nM NH-Fork DNA+10 nM SMARCAL1+220 nM RAD52-OBD</td><td>3</td><td colspan="2">77±24.0 (4.8%)</td><td>167±32 (8.6%)</td><td colspan="2">491±51 (86.7%)</td><td colspan="2"></td></tr>
<tr><td>12. 10 nM NH-Fork DNA+10 nM SMARCAL1+330 nM RAD52-OBD</td><td>3</td><td colspan="2">74±24.1 (13.2%)</td><td>167±30 (8.0%)</td><td colspan="2">495±53 (78.8 %)</td><td colspan="2"></td></tr>
</table>

* Replication fork bound to both a single ring of RAD52 and SMARCAL1

The data in this table are representative of three independent experiments. All data were analyzed using DiscoverMP software package. The molecular weights correspond to the average (± standard deviation) of respective Gaussians peaks. The percentage of the counts in each peak is shown in the parenthesis.

**Supplemental Figure S14**

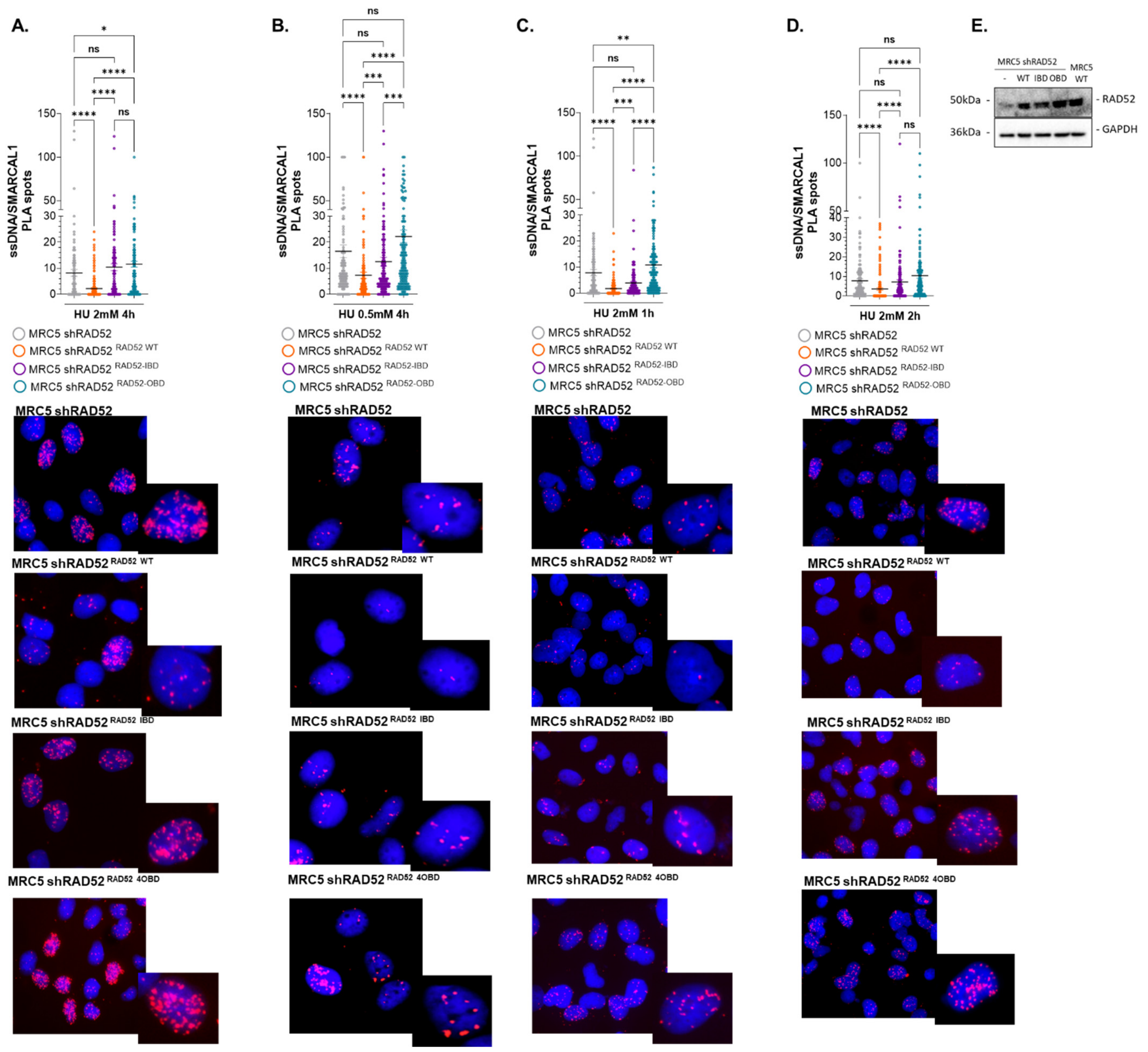


**Competition between RAD52 and SMARCAL1 is observed at different time points after induction of replication stress and at different HU concentrations. A.** The fork:SMARCAL1 experiment from **Fig. 4G. – J.** Analysis of SMARCAL1-parental ssDNA interaction by PLA. RAD52 knockdown cells, complemented with the indicated RAD52 mutants were treated with 100 μM IdU for 20 hours, released for 2 hours in fresh medium and exposed to HU at different time-points (**A, C, D**) or at low HU dose (**B**). The PLA reaction was carried out using antibodies against the SMARCAL1 protein and IdU. Representative images are shown. Magnification of one nucleus is presented in the inset. Graphs report the quantification of the number of PLA spot per nucleus in each condition represented by a different form of RAD52 (ns = not significant; **P<0.1; ***P < 0.001; ****P < 0.0001; Kruskal-Wallis test). **E.** WB showing the similar expression of each RAD52 isoform in the PLA experiments.

**Supplemental Figure S15**

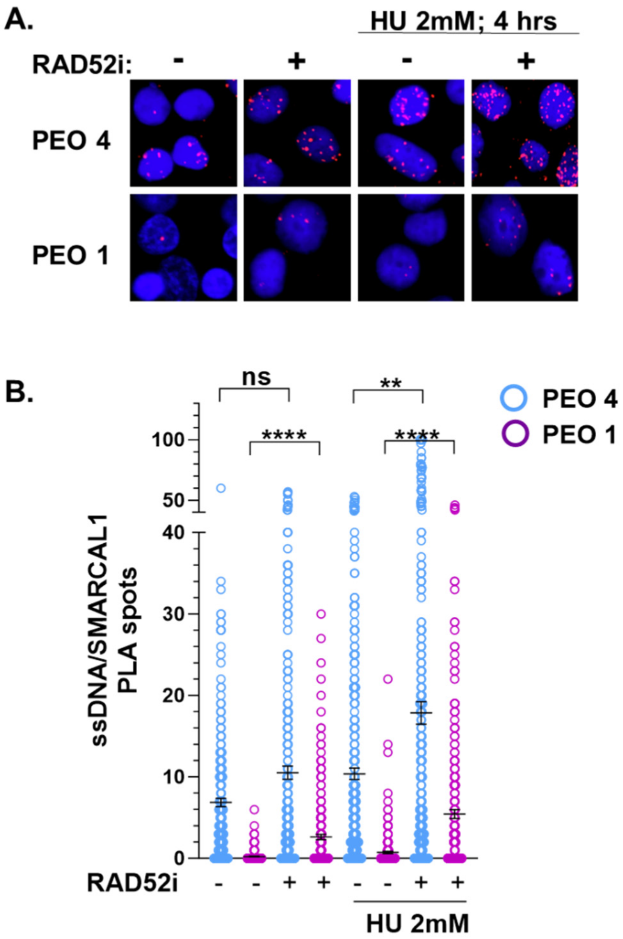


**Competition between RAD52 and SMARCAL1 is observed also in a BRCA2-mutated background. A.** Analysis of SMARCAL1-parental ssDNA interaction by PLA. The BRCA2-mutated cancer cell line PEO1 and its spontaneous revertant, BRCA2-proficient, clone PEO4 were treated with 100 μM IdU for 20 hours, released for 2 hours in fresh medium and exposed to HU. The PLA reaction was carried out using antibodies against the SMARCAL1 protein and IdU. Representative images are shown. **B.** Quantification of the number of PLA spot per nucleus. (ns = not significant; **P<0.1; **P < 0.01; ****P < 0.0001; Kruskal-Wallis test).

**Supplemental Figure S16:**

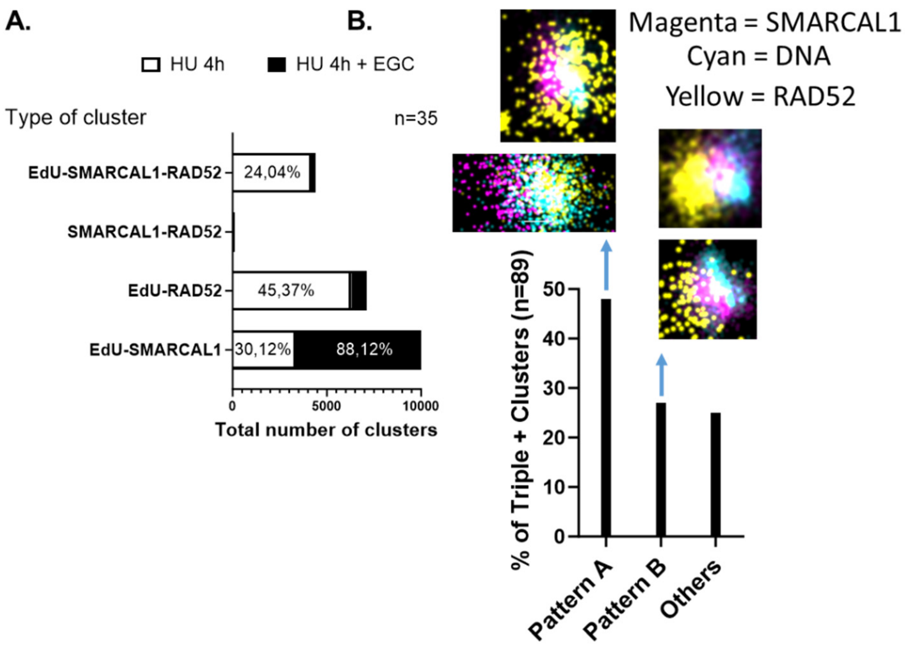


**Analysis of the RAD52 and SMARCAL1 localization by super-resolution microscopy. A.** Quantification of EdU-RAD52-SMARCAL1 interaction by dSTORM. U2OS cells were treated as indicated on top. The graph shows the total number of the indicated localization clusters. Numbers indicate the percentage of co-localization events. N denotes the number of images from two independent sets. **B**. Topological assessment of the distribution of the indicated signal in the triple-positive clusters. The graph shows the percentage of each pattern in the clusters (n=89). Representative dSTORM images of each patter are shown on top of the relevant histogram bar. Scale bar = 100nm. All the values above are presented as means ± SE (ns = not significant; *P < 0.1; **P<0.1; ***P < 0.001; ****P < 0.0001; Kruskal-Wallis test)

### Supplemental Figure S17

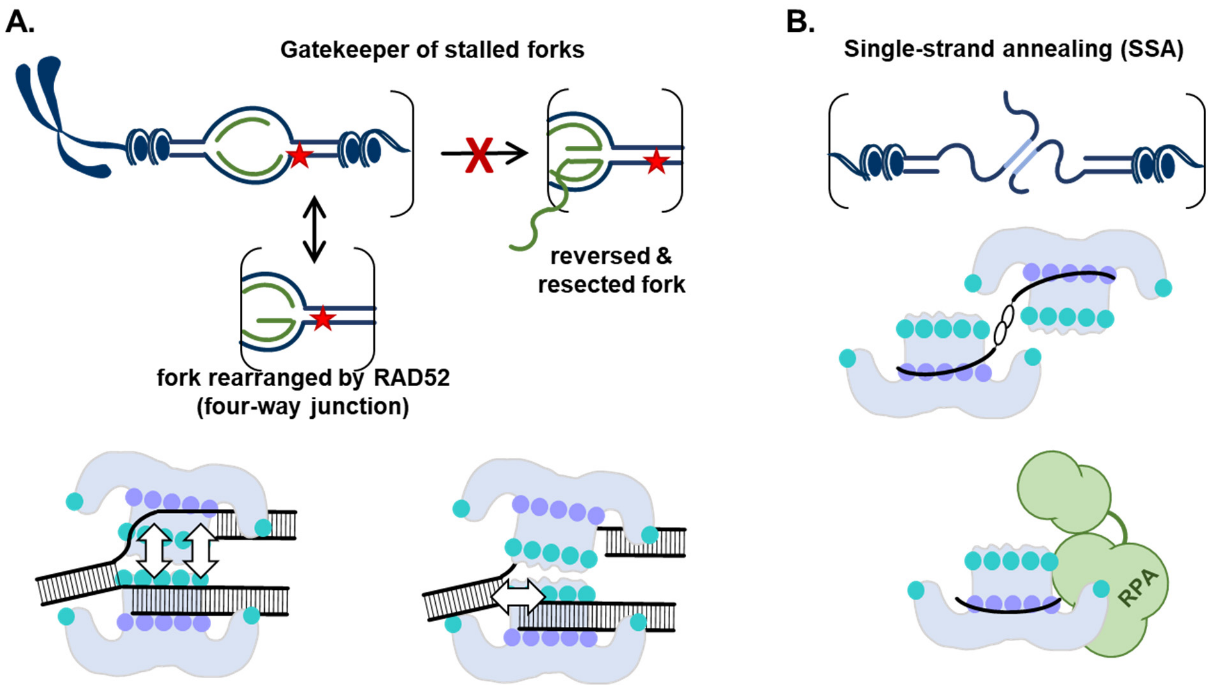


**Proposed change in the RAD52-DNA architecture during different cellular functions. A.** As a gatekeeper of stalled DNA replication forks[2], RAD52 forms a double-ring spool-like structure (**Fig. 2E.**) that incorporates all three arms of the replication fork and rearranges the fork through by exchanging the homologous DNA strands (**Fig 1C-H**, **Fig.3**, **Supplemental Figure S12**). Fork remodeling is distinct from that mediated bySMARCAL1 and other fork remodeling motors as it restricted to the gap region and does not continue into four-way junction branch migration. The double-ring structure of RAD52 directly competes with SMARCAL1 for the fork binding *in vitro* and in cells (**Fig. 4**) and also creates a four-way junction substrate that SMARCAL1 can restore back to the replication fork[1] (and **Supplemental Figure S3 A-E**.), but is inefficient in further reversal (**Supplemental Figure S3 F-J**.). A spool-like architecture of the RAD52-fork complex is important for the efficient fork engagement and the rearrangement. **B**. In contrast to the requirement of the gatekeeper function on the outer DNA binding site, single strand annealing by RAD52 is inhibited by the dsDNA binding[3]. High resolution X-ray crystallography of a complex between The N-terminal domain of RAD52 and DNA suggested that ssDNA annealing may involve relocation of the ssDNA from the narrow inner DNA binding sites of two RAD52 rings to the top of the ring where homology can be sampled[4]. Recent cryo-EM structures of the RAD52-ssDNA and RAD52-ssDNA-RPA complexes posed for strand annealing showed a similar placement of the ssDNA in the inner DNA binding site but with a broken decameric ring of RAD52 which accommodates both ssDNA and RPA[5].
